# Structural unification of diverse membrane-bound acyltransferases reveals a conserved fold that defines the Transmembrane Acyl Transferase (TmAT) superfamily

**DOI:** 10.1101/2024.11.11.622933

**Authors:** Bethan E. Kinniment-Williams, Vytaute Jurgeleviciute, Reyme Herman, James N. Blaza, Marjan W. van der Woude, Gavin H. Thomas

**Affiliations:** Hull York Medical School, University of York, York, UK; York Biomedical Research Institute, University of York, York, UK; Department of Biology, University of York, York, UK; York Structural Biology Laboratory, Department of Chemistry, University of York, York, UK

## Abstract

The movement of acyl groups across biological membranes is essential for many cellular processes. One major family of proteins catalysing this reaction are the acyl transferase family 3 (AT3) proteins, which form a pore to allow acyl-CoA to penetrate the membrane for transfer onto an extracytosolic acceptor molecule. Recent structures of the sequence-unrelated human heparan-α-glucosaminide *N*-acetyltransferase (HGSNAT) support a similar transmembrane acyl-group transfer mechanism. Here we demonstrate that both protein families contain a conserved 10-transmembrane helical fold with high structural and detectable sequence conservation around the acyl-CoA pore, supporting the previously proposed Transmembrane Acyl Transferase (TmAT) protein superfamily. In addition, we identify TmAT proteins, including the human Golgi sialate-O-acetyltransferase (CASD1), the human/fungal PIG-W/GWT1 enzymes and the bacterial vancomycin resistance protein VanTG, where the TmAT domain’s function has been largely unrecognised. We conclude that the TmAT fold represents an ancient architecture for transmembrane acyl-group transfer with important roles in the dynamic modification of glycans in diverse processes across the three domains of life.

---

The modification of sugars via O-acylation is used across biology to alter the structure and properties of glycans^1–3^, and impacts processes ranging from glycan degradation in the human lysosome^4–6^ to evasion of immune and viral attack on fungal and bacterial pathogens^7–10^. While many acylations occur in the cytosol using soluble enzymes that have access to acyl-CoA pools as acyl-donors^11^, there are instances where the sugar is not modified until it has been moved outside of the cytosol into another cellular compartment or surface of the cell^12,13^. Members of the Acyltransferase-3 (AT3)/Putative Acetyl-CoA Transporter (ATAT) family (TCDB 9.B.97) (InterPro IPR002656) of membrane bound proteins facilitate the extra-cytosolic modification of some of these sugars^2^. For example, O-acetylation of the oligopolysaccharide (OPS) in *Shigella flexneri* is catalysed by AT3 proteins, including OacB and Oac, resulting in a serotype conversion which is a recognised virulence determinant^14–17^. In *Staphylococcus aureus* and *S. epidermidis* IcaC is important for the development of poly-*N*-acetyl-glucosamine (PNAG) derived biofilms and has been suggested to O-succinylate the polysaccharide^18^. While both OacB and IcaC are thought to be standalone AT3 proteins, some members of the AT3 family are fused to a second soluble domain. A commonly identified fusion involves a SGNH hydrolase domain which is believed to participate in the transfer of the acyl-group from the AT3 domain to the extra-cytoplasmic carbohydrate acceptor^1,2,13,19^. An example of an AT3-SGNH fusion is the widely found peptidoglycan O-acetylase, OatA, again found in *S. aureus*. The O-acetylation of the GlcNAc sugar within peptidoglycan occurs after it has been exported out of the cytosol and confers the important phenotype to the cell of resistance to host lysozyme^12,13^. We characterised another example of an AT3-SGNH fusion in the form of the Salmonella LPS O-acetylase OafB^1^ which confers resistance to bacteriophage infection. Using computational methods, we identified a stable 10 transmembrane helix (TMH) core structure which contains a pore that accommodates the acetyl-CoA (AcCoA) donor^1^. Fused AT3 proteins are also found across taxa including eukaryotic proteins such as the *Drosophila melanogaster* Drop Dead protein (DRD)^20^ and the *Caenorhabditis elegans* nose fluoxetine resistance proteins^21^. Unlike their bacterial counterparts, these proteins have an NRF domain fused to the N-terminus of the AT3 domain.

The recent structures of the human membrane-bound heparan-α-glucosaminide *N*-acetyltransferase (HGSNAT) protein^4–6^, provide exciting and important information about intramembrane transfer of acyl-groups onto sugars in a protein that has been considered from sequence analysis to be unrelated to the AT3 family of acyltransferases. HGSNAT acetylates heparan sulfate for its subsequent breakdown in the lysosome and loss of function leads to the accumulation of heparan sulfate in the lysosome, causing diseases such as Sanfilippo syndrome^22^. In what appears to be a strikingly similar process to OafB, the HGSNAT protein binds acetyl-CoA on the cytosolic face, with the protein creating a pore for it to insert into the membrane and present the acyl-group to a catalytic site on the opposite side for movement onto an extra-cytosolic sugar. The HGSNAT protein also contains a second domain (called α-HGSNAT), although this luminal domain is structurally distinct from OafB’s SGNH domain and is cleaved after synthesis but stays associated with the 10 TMH membrane domain (called β-HGSNAT) through a single TMH. In two of these recent publications^4,5^, the AT3 proteins are not mentioned as a related family of membrane-bound acyltransferase. Additionally, Xu *et al*.^*4*^ report finding no structural matches to HGSNAT in their DALI search of the PDB due to a lack of solved structures; However, their search did not extend to the AlphaFold database which includes predicted structures.

Despite Xu *et al*.^4^ and Zhao *et al*.^5^ finding no structural similarities to HGSNAT, the Transporter Classification Database (TCDB) places OafB and HGSNAT into two different families (9.B.67 AT3 and 9.B.169 HGSNAT/YeiB, respectively), but importantly groups them into the same superfamily which is known as the Transmembrane Acyl Transferase (TmAT) superfamily^23^. The third paper by Navratna *et al*.^*6*^ recognises the TmAT superfamily, noting their structure of HGSNAT as the first within this superfamily. They highlight the similar architecture between the transmembrane domains of an AT3 OafB and HGSNAT but ultimately conclude that the structures are not homologous. In this analysis, we present the first evidence that there is clear structural homology across the TmAT superfamily, including homology between OafB and HGSNAT. We also undertake a detailed analysis of a larger grouping of membrane-bound proteins from several distinct families, enabling us to refine the membership of the Acyl_transf_3 Clan in Interpro to include important unrecognised members, enabling us to define conserved sequence characteristics of the TmAT fold, which cluster around their unique membrane-embedded acyl-CoA binding site. As this analysis confirms at a structural level the proposed TmAT superfamily^23^ we suggest this name is adopted more widely to represent proteins with this fold found in biology.

## Analysis

### Proteins in the TmAT superfamily contain a core conserved 10 TMH fold

To assess whether the membrane domain architecture is homologous within the proposed TmAT superfamily we first compared the structural similarity of the best structurally characterised AT3 protein, OafB, to the human HGSNAT using DALI tools^24,25^. As both proteins contain additional non-homologous domains (***Fig*.*1A***), they were excluded from the analysis. A common structure is observed for both membrane domains (***Fig*.*1B-D***), which includes core helices 1-4 of OafB (2-5 of HGSNAT) which are involved in acyl-CoA binding, the scaffolding helices 5-8 of OafB (6-9 of HGSNAT), which sit ‘behind’ the core helices, and the final two helices, 9-10 of OafB (10-11 of HGSNAT) the first of which is an unusually bent helix that closes one side of the acyl-CoA binding site^4–6^. The Z score and RMSD between the 10TM membrane domains of OafB and HGSNAT is 13.2 and 5.0 Å with 73% coverage, supporting the hypothesis that these proteins are structurally related. To test this idea more rigorously, we widened the analysis to 16 additional members from each of the two TCDB-defined families that contain OafB and HGSNAT (9.B.67 AT3 and 9.B.169 HGSNAT/YeiB) with representatives from bacteria, archaea and eukaryotes. A DALI “all-against-all” comparison was used to demonstrate conservation of the fold across all examined members (***Supplementary Fig*.*1-4, Supplementary Table 1-2***).

**Fig. 1.**
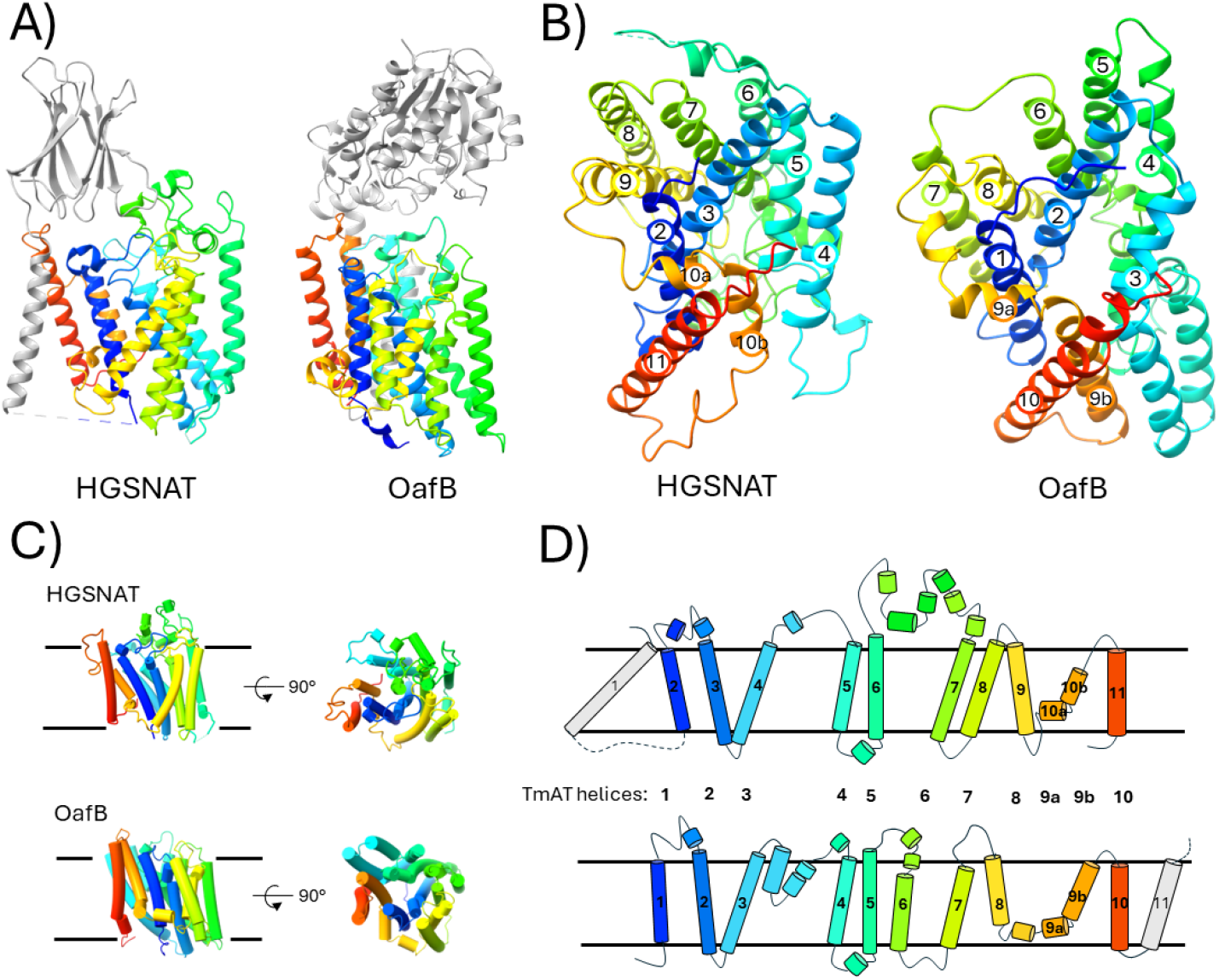
Conservation of a core 10-TMH TmAT protein fold. A) Side view of HGSNAT and OafB showing their additional fused domains and helices (grey). B) Cytosolic view of the core membrane fold from HGSNAT and OafB. C) and D) Cartoon representation of the helices of HGSNAT and OafB illustrating common elements and structural organisation where dotted lines represent additional structures. Proteins are all coloured with rainbow colouring (N-terminus to C-terminus). HGSNAT uses PDB code 8JKV and OafB the AlphaFold model for OafB (UniProt ID A0A0H2WM30), both are apo-states without acetyl-CoA bound.

### Human and fungal GWT1/PIGW proteins are members of the TmAT superfamily

Having established the structural similarity of the TmAT superfamily as defined by the TCBD, we also then critically assessed the structural similarities within the Pfam Acyl_transf_3 Clan (CL0316) that contains 9 Pfam families (***Supplementary Table 3***). Using our DALI-based searches we also discovered an additional Pfam family with the same fold, the GWT1/PIGW family of membrane-bound acyltransferases, that we included in this analysis. A DALI-based correspondence analysis of the structural similarity of these proteins strongly supports 7 of the 9 original families as being truly homologous, which increases to 8 with the inclusion of the GWT1/PIGW family (***Fig*.*2***). Two of the original families included in the clan, PF11318 (DUF3120) and PF12291 (DUF3623) are significantly shorter and have no shared structural organisation with the TmAT fold. The TraX family (PF05857) is the most divergent of the protein families that we consider to have the TmAT fold, lacking the characteristic broken 9th TMH and the 10th TMH. This divergence might be explained as this protein is known biologically to be required for acetylation of a protein substrate^26^, the bacterial F-pilin protein, so differs from the carbohydrate acceptors that are usually seen for TmAT proteins. In this analysis we also included control proteins from the MBOAT family of membrane-bound acyltransferases such as bacterial DltB and human Hedgehog acyltransferase (HHAT)^27,28^, which are clearly structurally distinct from the TmAT fold (***Fig*.*2***), as previously described^1^. Remarkably, during the preparation of this Analysis, the first structure of a GWT1/PIGW protein was solved^29^, confirming our hypothesis about its membership of the TmAT superfamily and the power of using AlphaFold/DALI-based searches for structural homology.

**Fig. 2.**
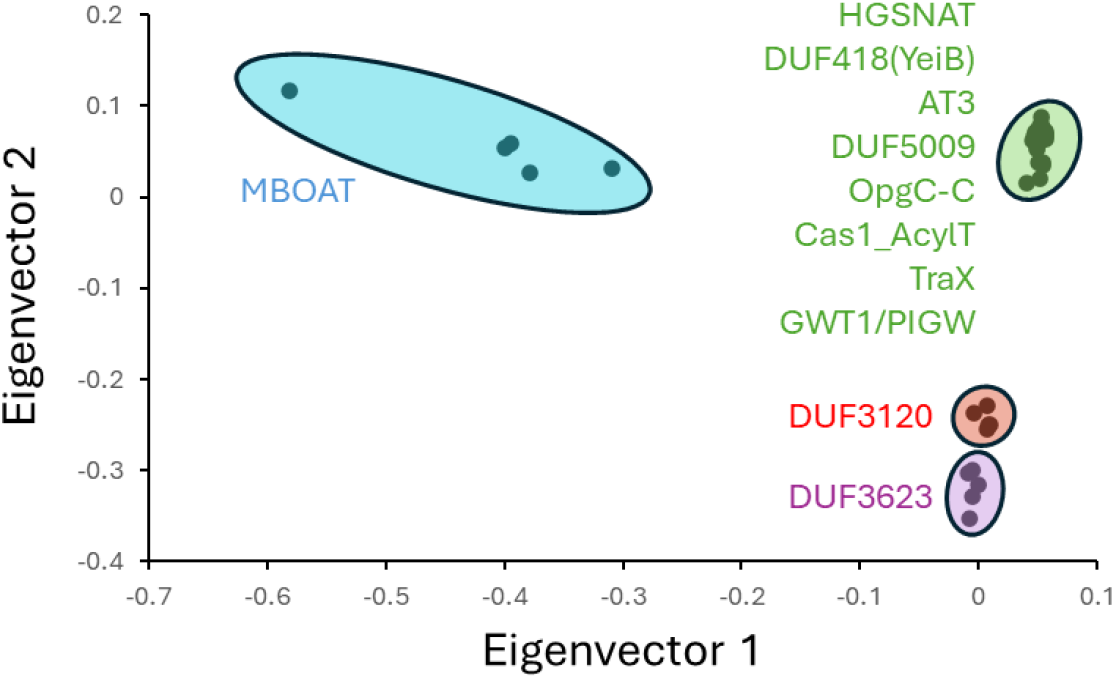
Structural analysis of the TmAT superfamily refines the membership of the Pfam Acyl_transf_3 Clan (CL0316). Correspondence analysis of predicted structures of representative proteins from the nine Pfam families that constitute the Pfam Acyl_transf_3 Clan (CL0316) (***Supplementary Table 3***), plus the GWT1/PIGW (PF06423) and MBOAT family (PF03062). MBOAT was included as an out-group of a structurally unrelated membrane-bound acyltransferase. This multidimensional scaling method uses eigenvectors to separate structurally different proteins on two different axes, so the most structurally similar proteins are positioned together. Five members of each were selected for structural correspondence analysis in DALI, with full-length protein representatives from bacteria, eukaryotic and archaea where possible (see ***Supplementary Fig*.*5-6 & Supplementary Table 4***). Note that three proteins that were previously annotated as DUF5009 have now been annotated as HGSNAT or are unlabelled (see Methods). Colours represent the different structural groups we define from the analysis.

### The most highly conserved sequence and structural elements of the TmAT superfamily form the acyl-CoA binding site

While we have confirmed the structural homology of the TmAT superfamily across its two constituent TCDB families and refined the Pfam Clan with the same fold, we next sought to use our structural alignments to uncover if there was any sequence conservation across these functionally diverse proteins (***Fig*.*3***). We then compare these findings to the limited number of studies where mutagenesis has been used to study the function of proteins in the TmAT superfamily. Within our alignments of the members of the superfamily (***Fig*.*3***), there were 7 positions that were highly conserved, with 5 occurring in TMH 1-3 and the others in TMH 6 and TMH 9, consistent with the most conserved structural features of the fold (***Supplementary Fig*.*7***). We refer to these residues as TMH#-Xxx, where TMH# is the TMH harbouring the residue and Xxx is the amino acid. The corresponding residues in HGSNAT and OafB are found in ***Supplementary Table 5***.

**Fig. 3.**
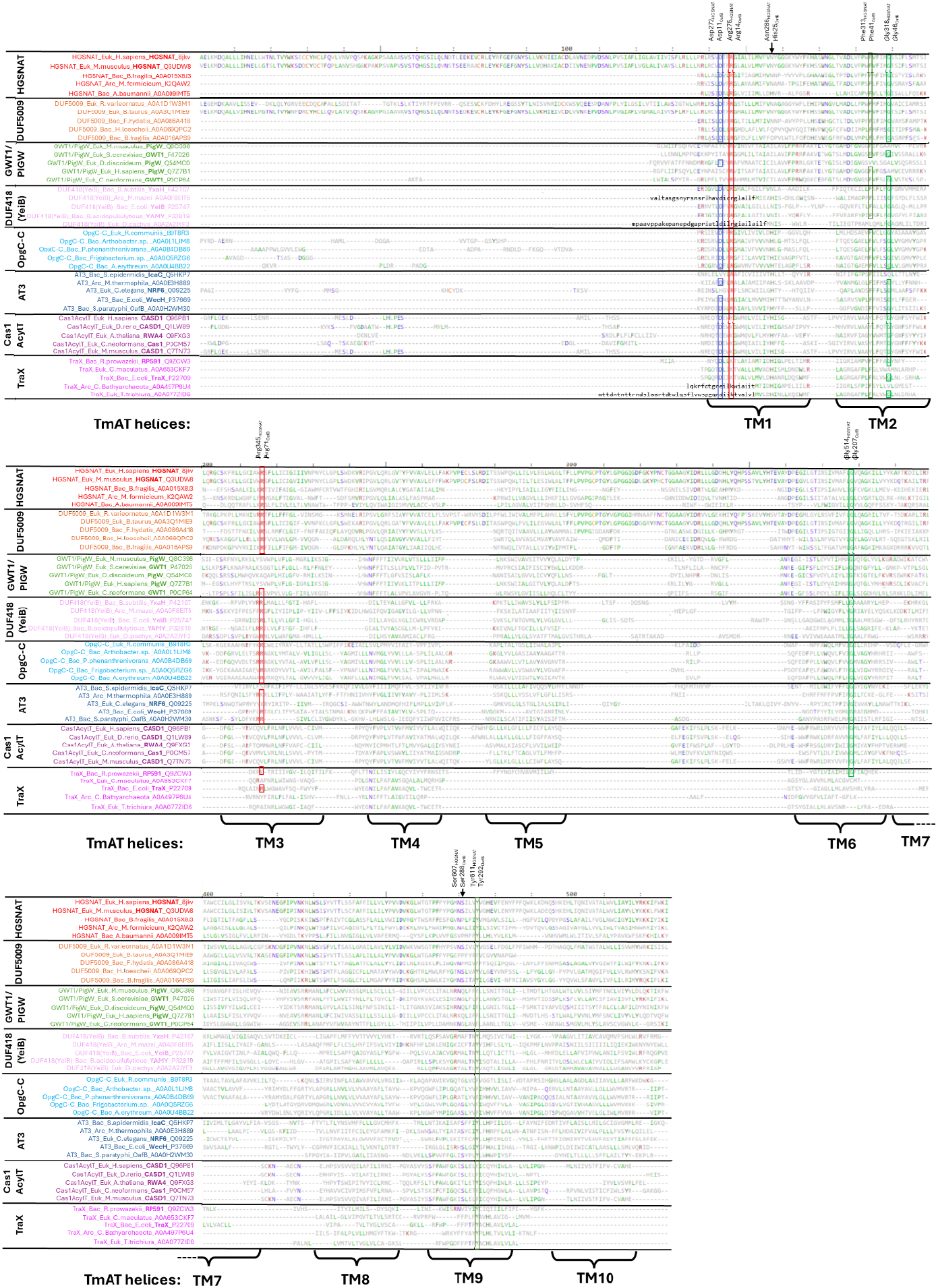
Structural alignment of the TmAT superfamily defines the core conserved residues. Structure-based alignment (compared to HGSNAT 8JKV) produced in DALI showing diverse representatives of all Pfam Acyl_transf_3 Clan (CL0316) members containing the TmAT domain (see ***Fig*.*2***) and the additional PigW/GWT1 family found by a DALI search. Regions with no structural/sequence homology are not expanded, so only structurally equivalent positions to HGSNAT (8JKV) are shown. For TraX A0A077ZID6, TraX A0A497P6U4, YeiB A0A2A2JYF3, and YeiB A0A0F8EIT5, these sequences were misannotated at the N-terminus and we manually searched for the most likely start codon (in black and lowercase). It is important to note that some of the proteins appear to not have a TM5 or TM7, but this is due to poor structural alignment with other TmAT members and not because they lack a TM5 or 7. Note that three proteins that were previously annotated as DUF5009 have now been annotated as HGSNAT or are unlabelled (see Methods). The most common amino acid at any given position is coloured according to aromaticity (dark green) (Phe, His, Trp, Tyr), positive charge (red) (Arg, Lys), negative charge (blue) (Glu, Asp), polar uncharged side chains (purple) (Ser, Thr, Gln, Asn), hydrophobic side chains that are not aromatic (Gly, Leu, Ile, Val, Pro, Met, Ala), and sulfur containing amino acid (orange) (Cys). Boxed residues are those that are conserved, or contain conservative substitutions (Phe?Tyr, Arg?Lys, Asp?Glu), across the alignment. Additional partially conserved residues discussed in the text are marked with an arrow. See ***Supplementary Fig*.*9*** for secondary structure alignment.

Working through the proteins, the TMH 1 contains a conserved DxxR motif with both residues positioned at the cytosolic side of the protein. While the Asp (TMH1-Asp) in this motif is essential for the function of HGSNAT, it appears to be dispensable for the function of the AT3 protein, OacB (Asp44)^4,14^ (mutated residues are shown in ***Supplementary Fig*.*8***). However, the Arg residue (TMH1-Arg) is essential for the function of HGSNAT, OafB and OacB (Arg47)^4,7,14^. It is of note that in two of the Pfam families included in the Acyl_transf_3 Clan, Cas1 and TraX, the Arg is usually replaced by a Lys. Moving into TMH2, there is a conserved FxxxxG motif, where the Phe (TMH2-Phe) has been shown to be important for the activity of HGSNAT and the AT3 protein, OatA (Phe52)^4,13^. However, the constituent conserved Gly (TMH2-Gly) was found to be non-essential in OatA ^13^, and no mutagenesis studies were conducted on this residue in HGSNAT. In AT3 proteins, the widely recognised TMH3 RxxR motif has been shown to be important for the function of OafB, OafA, Oac, and OacB^2,4,8,13–16^. The first Arg (TMH3-Arg) of this motif is conserved in most members of the TmAT superfamily and its mutation in HGSNAT causes a significant loss of activity^2,4,8,13–16^. An additional highly conserved Gly (TMH6-Gly) was identified in TMH6 and in HGSNAT, a missense mutation of this residue (G514E) is found in patients with Sanfilippo type C disease^7,30^. The final conserved residue is a Tyr in the bent TMH 9 (TMH9-Tyr). Although this position has not been investigated via mutation in HGSNAT, Tyr611 surrounds the beta-mercapto-ethylamine of acetyl-CoA^4^ and the equivalent residue in OatA has been shown to be important for protein function^13^. In AT3 proteins, this Tyr is part of the SxxxY motif in TMH 9^1^ which is also conserved in the human HGSNAT protein. However, the Ser (TMH9-Ser) is not conserved across all members of the superfamily, and the equivalent residue has been shown to be non-critical for the function of OatA (Ser310)^13^.

Given the newly identified sequence and structural features of the TmAT superfamily, we assessed how these most conserved elements might be involved in enzyme function. Strikingly, but perhaps not surprisingly, the most conserved elements of the TmAT proteins are involved in the binding of the conserved acyl-CoA donor molecule (***Fig*.*4***). Building on prior work on AT3 proteins, which predicted the OafB acyl-CoA binding site^1^, the new structures of HGSNAT^4–6^ provide the precise location of the acetyl-CoA binding site. This location is largely consistent with the location predicted by molecular dynamic simulations for OafB^1^.

**Fig. 4.**
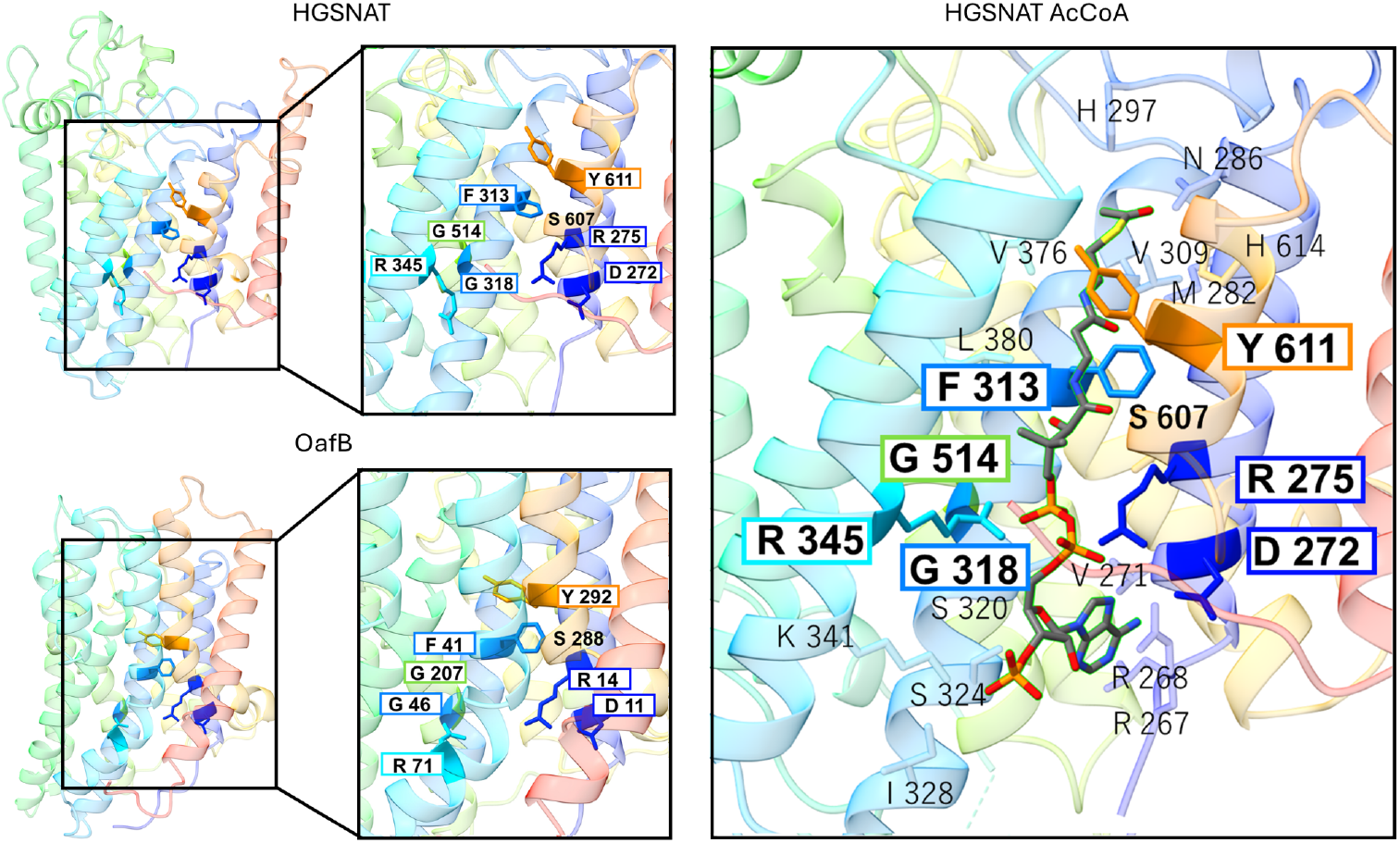
The conserved residues of the TmAT superfamily form the acyl-CoA binding site. Conserved residues in the TmAT superfamily superimposed on A) the 10TM domain of HGSNAT (8JKV) and B) the 10TM domain of OafB, illustrating their common arrangement and in C) with acetyl-CoA bound in HGSNAT (8JL1) illustrating their role in acyl-CoA recognition. The seven highly conserved residues are boxed and in bold text, with the additional partially conserved residue, Ser607 (TMH9-Ser) (see text) in bold, but unboxed. TMH1 is shaded in dark blue, TMH2 in blue, TMH3 in cyan, TMH6 in green, and TMH9 in orange. The residues shown in a smaller text represent the residues interacting with acetyl-CoA according to the PDB structure 8JKV^4^. Note that the F313 and R345 structurally reorient during acetyl-CoA binding^4^.

The conserved amino acids form what appears to be an equivalent binding site formed by the same helices in HGSNAT (TM2-5, TM10, TM11) and in AT3 (TM1-4, TM9, TM10) (***Fig*.*4***). Acetyl-CoA binds in an extended conformation to enable it to enter the protein from the cytosol and penetrate through the membrane to the enzyme’s catalytic site on the extra-cytosolic face (***Fig*.*4***). The conserved residues in the TmAT superfamily play key roles in the interactions of the protein with the full length of the acyl-CoA molecule in its binding site, with the majority interacting with the ‘common’ adenosine and the 4-phosphate pantothenic acid moiety of CoA found in all acyl-CoA substrates (***Fig*.*4***). These are on the cytosolic side and should be the most conserved elements given that our alignment includes TmAT superfamily members that use succinyl-CoA and other longer chain acyl-CoA as acyl-donors^18,29,31^, where the structure of the acyl-donor varies beyond the thioester bond. Interestingly the ‘family specific’ conserved residues are found to line the acyl-CoA binding site further towards the extracytosolic side (***Fig*.*4***). This includes an important Asn residue in HGSNAT further along TMH 1, which in AT3 proteins is found as a His residue in a family-specific DxxRx_10_H motif^1^. Both of these residues are important for function and have been proposed in some cases to be involved in catalysis following acyl-CoA binding^4,13^.

Taken together, these analyses of the shared structural and sequence conservation of proteins in the AT3 and YeiB/HGSNAT families entirely support the concept of them forming the TmAT superfamily and having a conserved evolutionary ancestry.

### Assessment of new members of the TmAT superfamily

Having established the power of comparing AlphaFold-derived structural models using DALI to establish the evolutionary basis of the TmAT superfamily, we looked deeper into the results to seek out further important proteins, that are either currently not classified into the two constituent families in the TCBD or are in Pfam families not currently included in the Acyl_transf_3 Clan (CL0316). Using the bacterial AT3 protein OafB as a query against the Human AlphaFold database in DALI, HGSNAT is returned as a strong match (Uniprot ID: Q68CP4, Z score: 13.1, RMSD: 4.7 with 37% coverage) in addition to two other interesting proteins, the CASD1 protein (Uniprot ID: Q96PB1, Z score: 17.1, RMSD: 4.5 with 34% coverage) and PIGW/GWT1 protein (Uniprot ID: Q7Z7B1, Z score: 12.1, RMSD: 4.6 with 48% coverage), both of which are discussed below.

#### PIGW/GWT1

The PIGW/GWT1 proteins we have mentioned previously and presented evidence that they sit within the TmAT superfamily (***Fig*.*2*** and ***Fig*.*3***), despite not being previously recognised as being in either the Pfam Acyl_transf_3 Clan (CL0316) or the TCDB TmAT superfamily are the PIGW/GWT1 proteins (PF06423, IPR009447). The new structure of the fungal GWT1 protein confirms this hypothesis^29^ (***Fig*.*5***) and closely resembles the AlphaFold model (***Suppplementary Fig*.*10-11***). Biologically these proteins are interesting TmAT members as they catalyse the addition of an acyl-group onto an inositol sugar acceptor in an early stage of the synthesis of glycosylphosphatidylinositol (GPI)^32,33^. Mutations in the human gene result in the loss of inositol acetylation, which reduces levels of GPI-anchored proteins and lead to West syndrome and hyperphosphatasia with mental retardation syndrome (HPMRS, also known as Mabry syndrome). In fungi the same protein, called GWT1, is required for the synthesis of their complex cell envelope and has been targeted in the development of novel anti-fungal^34^ and anti-parasitic drugs^35^. Our inclusion of these proteins in the TmAT superfamily has a functional consequence for these proteins, which is supported completely by the new structure. This relates to the spatial localisation of this reaction in the GPI biosynthesis pathway as the inositol acetylation step occurs before a mannosylation step^32^ that is catalysed on the luminal side of the ER membrane. Hence our structural model and the experimental structure now suggest that the immediately preceding inositol acetylation step would also likely occur on the luminal side of the ER membrane using cytosolic palmitoyl-CoA as the acyl-donor, rather than requiring there to be a pool of palmitoyl-CoA within the ER lumen as proposed by Sagane et al.^36^ GWT1 is also interesting as the acyl-group donor is much longer than other TmAT proteins (palmitoyl is C16) and the structure reveals that while the CoA part of the palmitoyl-CoA binds in a similar way to other TmAT proteins^29^, the longer fatty acid attached to the thiol group bends back into a broader channel in the protein, which is known to be able to accommodate acyl-CoA’s as short as C10 *in vitro*^*37*^, suggesting more flexibility, which is borne out by some sequence differences in the GWT1/PIG-W proteins compared to other TmAT superfamily members (***Fig*.*3***).

**Fig. 5.**
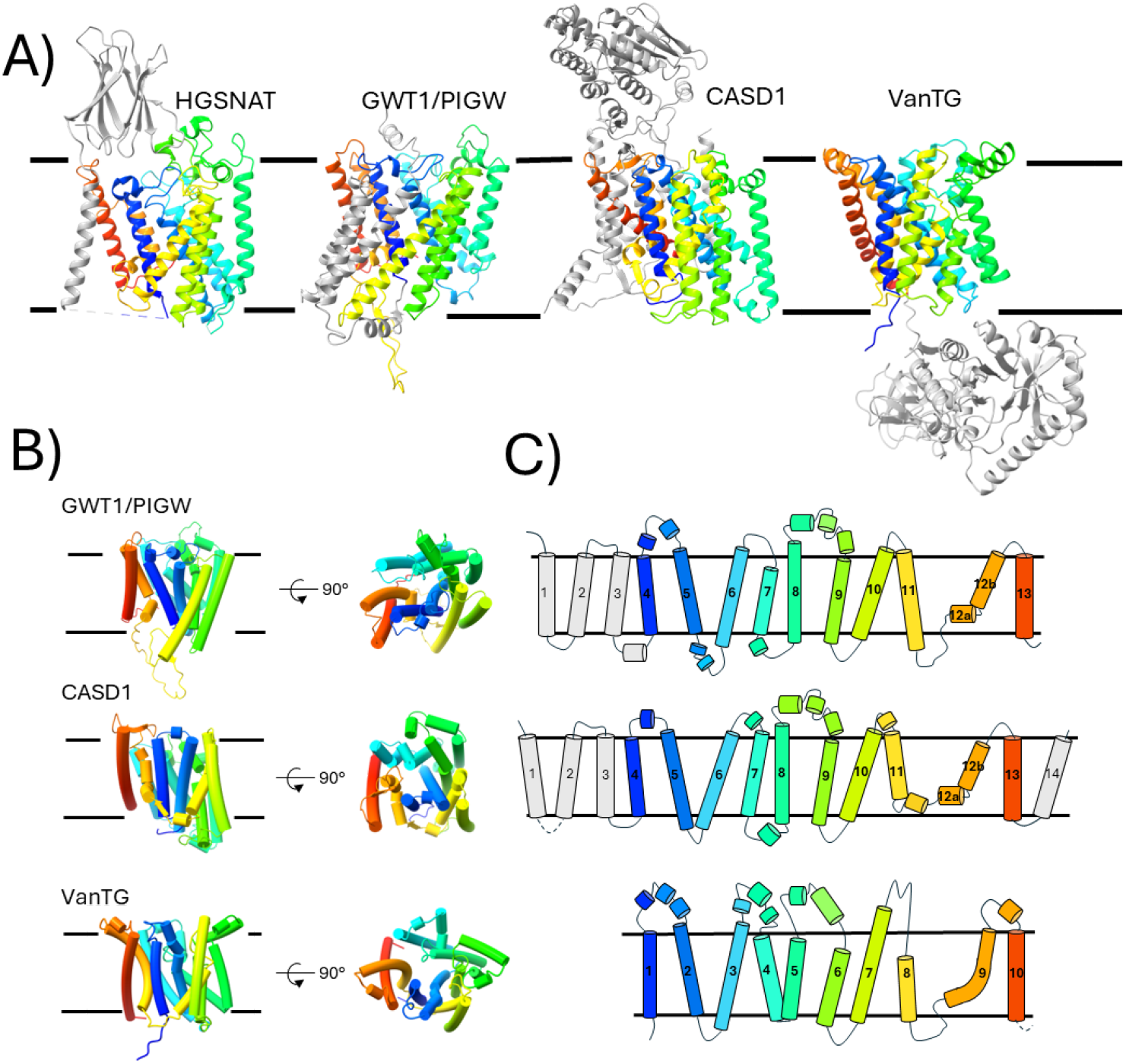
Presence of the core TmAT fold implies previously unrecognised additional biochemical functions in diverse biologically characterised proteins. A) Side view of HGSNAT, GWT1/PIGW, CASD1 and VanTG showing their additional fused domains and helices (grey). B) and C) Cartoon representation of the helices of HGSNAT, GWT1/PIGW, CASD1 and VanTG illustrating common elements and structural organisation where dotted lines represent additional structures. Proteins are all coloured with rainbow colouring (N-terminus to C-terminus). The following PDB or Alphafold IDs are used; HGSNAT (PDB ID: 8JKV), GWT1/PIGW (Uniprot ID: Q7Z7B1), CASD1 (Uniprot ID: Q96PB1), VanTG (Uniprot ID: Q186I3), all are apo-states without acyl-CoA bound.

#### CASD1

The human sialate O-acetyltransferase CASD1 (***Fig*.*5***) is an important Golgi protein that is essential for O-acetylation of sialic acids during the maturation of glycans and a potential target for preventing cancer progression^38^. While it has been previously noted to be a homologue of the *Cryptococcus neoformans* Cas1p protein that O-acetylates yeast glycans^39^, the significance of it being a two-domain protein that includes a TmAT domain has not been fully appreciated. Like the bacterial OafB protein, CASD1 also contains an SGNH transferase domain in the Golgi lumen. In OafB this domain catalyses the final step in the acetylation reaction, accepting the acetyl-group from the AT3 domain and adding it to the sugar residues in the LPS^1,2^. Studies on the human CASD1 protein have largely focussed on this SGNH domain^39^, which is N-terminal in CASD1, while the C-terminal TmAT domain has not been studied biochemically. The recognition of this second domain opens the possibility that acetyl-CoA does not have to be transported into the Golgi, as is currently thought^3^, as the acetyl group could be delivered to the luminal SGNH domain via the TmAT domain, much like in the bacterial OafB protein^1^. Much of the evidence for a Golgi acetyl-CoA transporter comes from elegant work by Varki & Diaz^40^, who took purified Golgi and followed the fate of acetyl-CoA radiolabelled on the acetate. Strikingly they found that most of this label (75-85%) was found on O-acetylated sialic acids in the Golgi, which supports the idea that there is a specific connection between the acetyl-CoA pool and the sialic acid acceptors in the two cellular compartments that could be mediated by the two domains of CASD1. This would ameliorate the need for the transport of acetyl-CoA into the Golgi, which might lead to its turnover by many enzymes and not specific labelling of sialic acids. Perhaps consistent with this, the acetyltransferase activity was lost when the Golgi membrane was disrupted suggesting delivery of the acetyl-CoA via the TmAT domain of CASD1 is required for efficient acetylation^41^.

#### VanTG

Finally, we wish to highlight VanT, an important protein implicated in the resistance of pathogenic bacteria to the clinically important antibiotic vancomycin. VanT (for example UniProt ID Q186I3) contains two domains, a known AT3 domain fused to a cytoplasmic serine racemase domain (***Fig*.*5***). As a known mechanism of resistance to vancomycin involves altering its binding site in the cell wall from containing a D-Ala-D-Ala structure with a D-Ala-D-Ser structure, the function of VanT has only considered its serine racemase domain, as part of the pathway to replace D-Ala with D-serine, made from L-Ser by VanT^42,43^. However, mutations that arise both in the lab and in clinical samples that lead to high-level vancomycin resistance fall into the AT3 portion of the *vanT* gene and not the racemase portion^44^. Hence, the AT3 domain itself clearly has an important function that has not been considered, but the new insights into this domain should facilitate future analyses.

The improved knowledge of acyl-CoA binding in the TmAT superfamily, combined with insights into catalysis from the HGSNAT structures, can also now be used to shed light on other AT3 protein family members which have important biological phenotypes but poor understanding of biochemical function. These include for example the NDG-4 protein from *Caenorhabditis elegans* that regulates lifespan^21,45^ and the HIF-1 Hypoxia-inducible factor inhibitor Rhy-1 from *C. elegans*^*46*^, which together are only 2 of the 63 AT3-containing proteins encoded in the *C. elegans* genome^46^. We also note that several fungal AT3-family proteins are encoded within biosynthetic clusters of various natural products, including the cholesterol lowering drug squalestatin S1^47^. Given that we already know that in *Streptomyces* sp. there are antibiotic biosynthesis clusters that contain AT3 proteins^2^ where they are known to catalyse the final acylation step in the biosynthesis step following export across the inner membrane^48^, we propose that these fungal proteins, such as *Phomopsis amygdali* Orf9 in the fusicoccin cluster^49^ and *Cochliobolus lunatus* Clz18 in the Zaragozic acid A cluster^50^, also have likely functions in natural product biosynthesis.

In conclusion, in this Analysis we demonstrated the power of structural comparisons to accurately assign members to the TmAT superfamily and distinguish subfamilies otherwise not observed at the sequence level (***Supplementary Fig. 12***). We identified important residues and a conserved 10TMH architecture that define this TmAT superfamily which allowed us to identify previously unrecognised proteins doing transmembrane acylation reactions of glycans and excluding non-structurally related families. This enables us to use the exciting and important data on acyltransferase mechanism from the recent HGSNAT work^4–6^ and GWT1 protein^29^ to infer similar mechanisms across a broad range of protein functions from archaea to man and significantly advances our understanding of many known AT3 and other TmAT superfamily proteins.

