## Supplementary Information & Methods for "Structural unification of diverse membrane-bound acyltransferases reveals a conserved fold that defines the Transmembrane Acyl Transferase (TmAT) superfamily"

### Methods

#### Bioinformatics methods for DALI all against all analysis.

The DALI [1,2] all against all tool was used to compare the 32 proteins included in the analysis of the TCDB[3] defined TmAT superfamily. This tool was also used to compare the 55 proteins from the nine families in the Pfam Acyl_transf_3 Clan (CL0316), the GWT1/PIG-W family and the MBOAT superfamily which helped to refine the TmAT superfamily classification. This approach was used to generate the heat maps, multiple sequence alignments, structural alignments, structural dendrograms, correspondence analyses, and provided RMSD values and percentage sequence identity.

#### Selection method for proteins in the two families in the TmAT superfamily represented in the TCDB

For DALI all against all analysis, we selected 16 proteins using Uniprot (accessed between 08/24-09/24), or PDB if available, from the Acyl_transf_3 (PF01757), and another 16 from DUF418 (PF04235)/HGSNAT-cat (PF07768). 6 eukaryotic members, 5 bacterial members and 5 archaeal members were included for each member of the TCDB defined TmAT superfamily (32 proteins were included in this analysis in total). Exact Uniprot ID’s / PDB ID’s, along with the domain and species, can be seen for these 32 proteins in ***Supplementary Fig.3*** (Structural dendrogram). The names follow the format as follows: ProteinFamily_Domain_Species_PROTEINNAME_UniprotID/PDBID. Protein names are in bold in ***Supplementary Fig.3*** unless they have yet to be named/are unreviewed. To focus the alignment on the core membrane helices, additional helices at either end and addition N- or C-terminal globular domains were removed (for example, the □-HGSNAT domain was excluded from HGSNAT, and C-terminal SGNH domain plus the linker was excluded from OafB). To see the exact truncations that were used in the analysis, see ***Supplementary Table 1.*** Proteins were aligned in CCP4MG [4] via Secondary Structure Matching (SSM) to human HGSNAT (PDB ID: 8JKV). Identifying the overlap between the TmAT 10TM in HGSNAT and other TmAT superfamily members allowed us to identify the 10TM TmAT fold in the other TmAT superfamily members. These superpositions were then exported as PDB files and put into ChimeraX1.8 where they were cut down to the appropriate 10TM helices.

#### Selection method for proteins from the the Pfam Acyl_transf_3 Clan (CL0316), GWT1/PIG-W family and MBOAT superfamily

For the DALI all against all analysis, the following selection method was used for the proteins that were included. Five proteins using Uniprot (accessed between 08/24-09/24), or PDB if available, were selected from following families; Acyl_transf_3 (PF01757), DUF418 (PF04235), HGSNAT-cat (PF07768), TraX (PF05857), Cas1_AcylT (PF07779), OpgC-C (PF10129), DUF3120 (PF11318), DUF3623 (PF12291), DUF5009 (PF16401), GWT1/PIGW (PF06423), and the MBOAT (PF03062) which served as our negative control (55 proteins were included in this analysis in total). Additional domains were not removed for this analysis. It is important to note that three of the proteins (Uniprot ID: A0A3Q1MIE9, A0A1D1W3M1, and A0A069QPC2) within this analysis were previously annotated as containing DUF5009 domains, but have since been reclassified. When Uniprot was accessed between 08/24 and 09/24, these proteins were annotated as having a DUF5009 domain. However, as of 04/11/24, A0A3Q1MIE9 and A0A069QPC2 are now annotated as HGSNAT, and the domain of A0A1D1W3M1 that was annotated as DUF5009 is no longer labelled within the Uniprot database. Within the AlphaFold database, two of these three proteins (Uniprot ID: A0A069QPC2 and A0A1D1W3M1) are still labelled as DUF5009, while A0A3Q1MIE9 is labelled as HGSNAT. Given that the structure of HGSNAT was recently solved and the TCDB's classification of the 9.B.97 family (HGSNAT/YeiB) as including DUF5009 proteins, it is unsurprising that the closely related DUF5009 has been reannotated in some cases. Exact Uniprot ID's / PDB ID's, along with the domain and species, can be seen for these 55 proteins in ***Supplementary Fig.6*** (Structural dendrogram). The names follow the format as follows: ProteinFamily_Domain_Species_**PROTEINNAME**_Uniprot ID's/PDB ID's. Protein names are in bold in ***Supplementary Fig.6*** unless they have yet to be named/are unreviewed. For this analysis we selected a mixture of bacterial, eukaryotic and archaeal members where possible. However, not all families included examples from all domains, so other domain protein members were used to make it up to 5. Generally, we also selected proteins that had been reviewed within Uniprot because they were more likely to have better data/more confident AlphaFold structures.

Multiple sequence analysis (MSA)

For multiple sequence alignments, all sequences were compared to the sequence of HGSNAT (PDB: 8JKV). Sequences are arranged by the family they belong to. Note that DUF3623, DUF3120 and MBOAT were excluded from the MSA analysis as they are not part of our defined TmAT superfamily. In the alignment of the revised Pfam Clan, if an amino acid was in more than or equal to 27/45 at a position (60% of TmAT superfamily) it was boxed. Amino acids were grouped together as (Phe, Tyr, His, Trp), (Ser, Thr, Gln, Asn), (Arg, Lys), (Glu, Asp), (Cys), (Gly), (Met), (Val), (Iso), (Leu), (Ala), (Pro). In the alignment for the TCDB-defined sequences, if an amino acid was in more than or equal to 27/32 at a position (84%), it was boxed. Amino acids were grouped together as (Phe, Tyr, His, Trp), (Ser, Thr, Gln, Asn), (Arg, Lys), (Glu, Asp), (Cys), (Gly), (Met), (Val), (Iso), (Leu), (Ala), (Pro).

DALI database search

The bacterial AT3 protein OafB (UniProt ID: A0A0H2WM30) was used as a query in the DALI search against the Human AlphaFold database. Strong matches to OafB were identified by a Z-score greater than 10.

Maximum likelihood phylogenetic tree

The amino acid sequences of the known or predicted membrane components (Supplementary Figure X) of the relevant proteins were aligned using MUSCLE 5.1 [5] and the maximum-likelihood phylogenetic trees was calculated using PhyML [6] using the Blosum62 substitution model and approximate likelihood ratio tests (aLRT). The resulting trees were visualised on Geneious Prime 2024.0.7 (https://www.geneious.com).

### Supplementary Figures and Tables

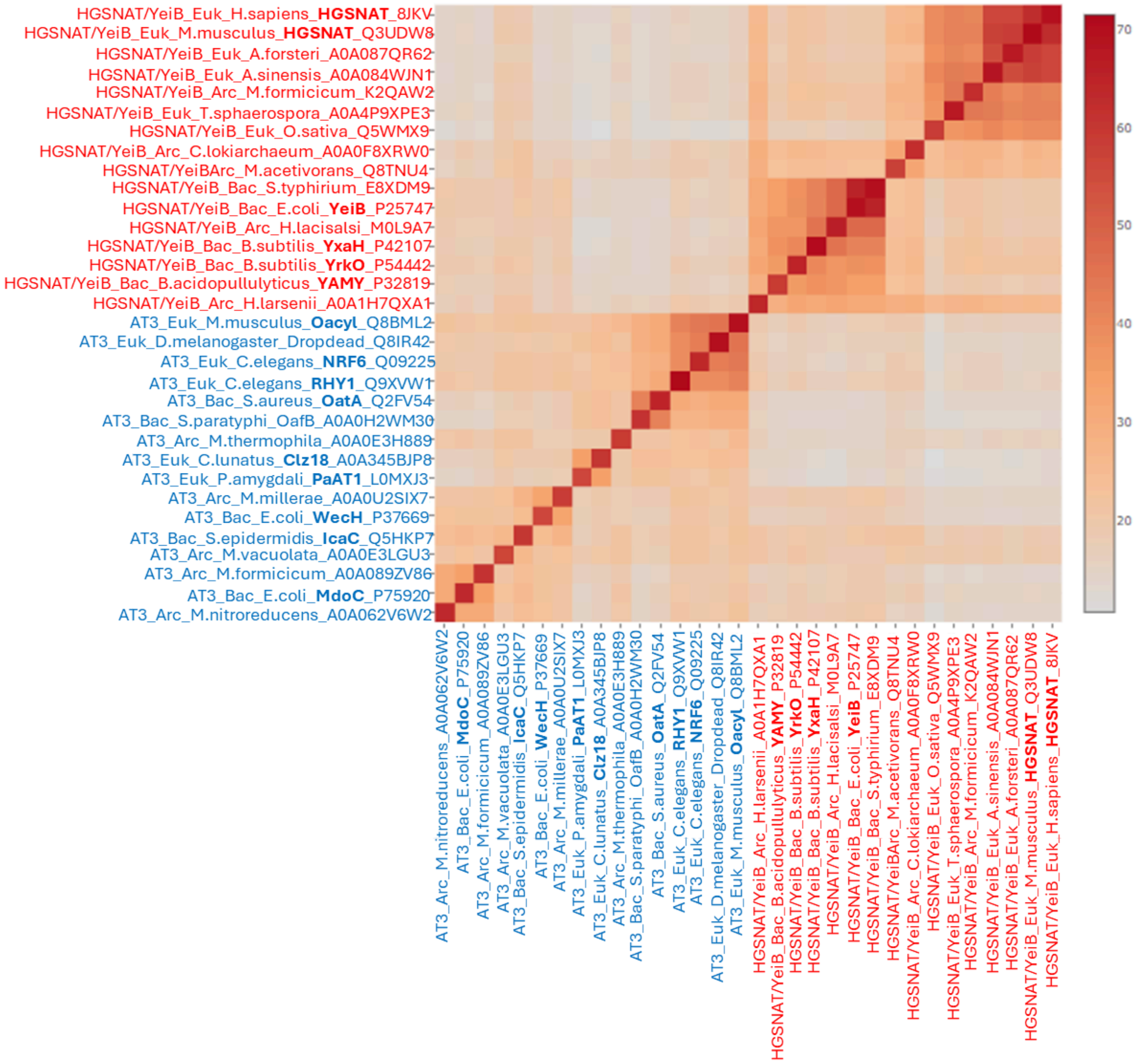


**Supplementary Fig.1. Heat map of Z-scores of a DALI all against all alignment of select members of the two TDBC-defined families (AT3 and HGSNAT/YeiB) within the TmAT superfamily**. Z scores of 16 members with representative members from archaea (Arc), bacteria (Bac) and eukaryota (Euk) are displayed. AT3 members are coloured in blue, while HGSNAT/YeiB members are coloured in red. The Z-score between two proteins is useful in determining homology: a Z-score below 2 suggests no significant similarity, while a score between 8 and 20 indicates a medium likelihood of homology and over 20, a high likelihood[7]. As expected, AT3 proteins show high Z-scores (13.7-45.3) when compared to each other, and HGSNAT/YeiB family proteins also display high Z-scores within their group (16.0-62.6). The protein members of the AT3 and HGSNAT/YeiB families appear to be at least distantly related, with all Z-scores being 10.5 or higher, confirming significant structural similarity across the TCDB-defined TmAT superfamily.

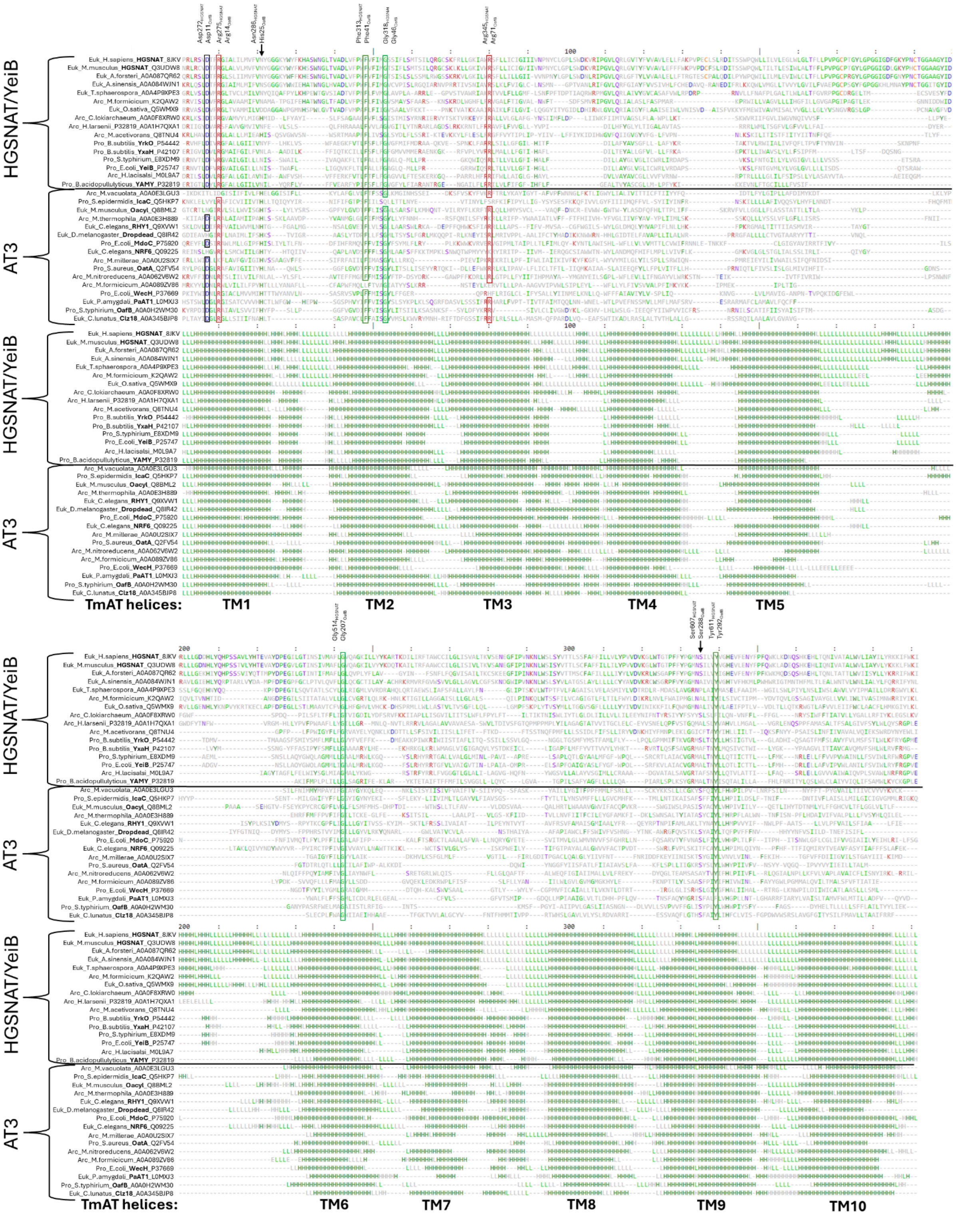


**Supplementary Fig.2. Structure-based sequence alignment of select AT3 and HGSNAT/YeiB family members.** The alignments were produced using DALI and HGSNAT (PDB: 8JKV) as the reference structure. Regions lacking any sequence/structural similarity are not expanded, so only structurally equivalent positions to HGSNAT (PDB: 8JKV) are shown. The most common amino acid at any given position is coloured according to aromaticity (dark green)(Phe, His, Trp, Tyr), positive charge (red) (Arg, Lys), negative charge (blue) (Glu, Asp), polar uncharged side chains (purple) (Ser, Thr, Gln, Asn), hydrophobic side chains that are not aromatic (Gly, Leu, Ile, Val, Pro, Met, Ala), and sulfur-containing amino acid (orange) (Cys). Boxed residues are conserved or exhibit conservative substitutions (Phe↔Tyr, Arg↔Lys, Asp↔Glu) across the alignment. Additional partially conserved residues discussed in the text are marked with an arrow. Protein secondary structure is annotated as H - Helix, L - Coil, and E -Sheet.

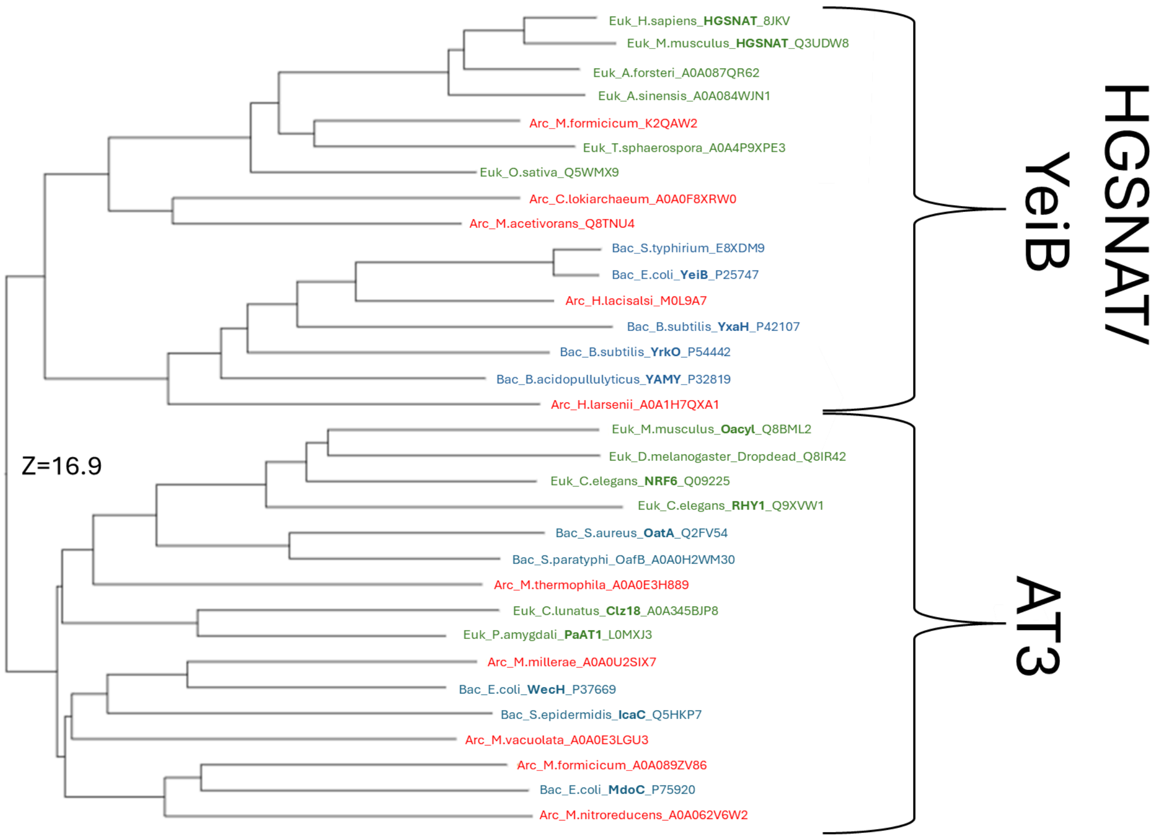


**Supplementary Fig.3. Structural dendrogram of select AT3 and HGSNAT/YeiB family members.** The structural dendrogram shows that the standalone AT3 and HGSNAT/YeiB families form two separate clades. Representative members from archaea (Arc), bacteria (Bac) and eukaryota (Euk) are coloured in red, blue and green respectively.

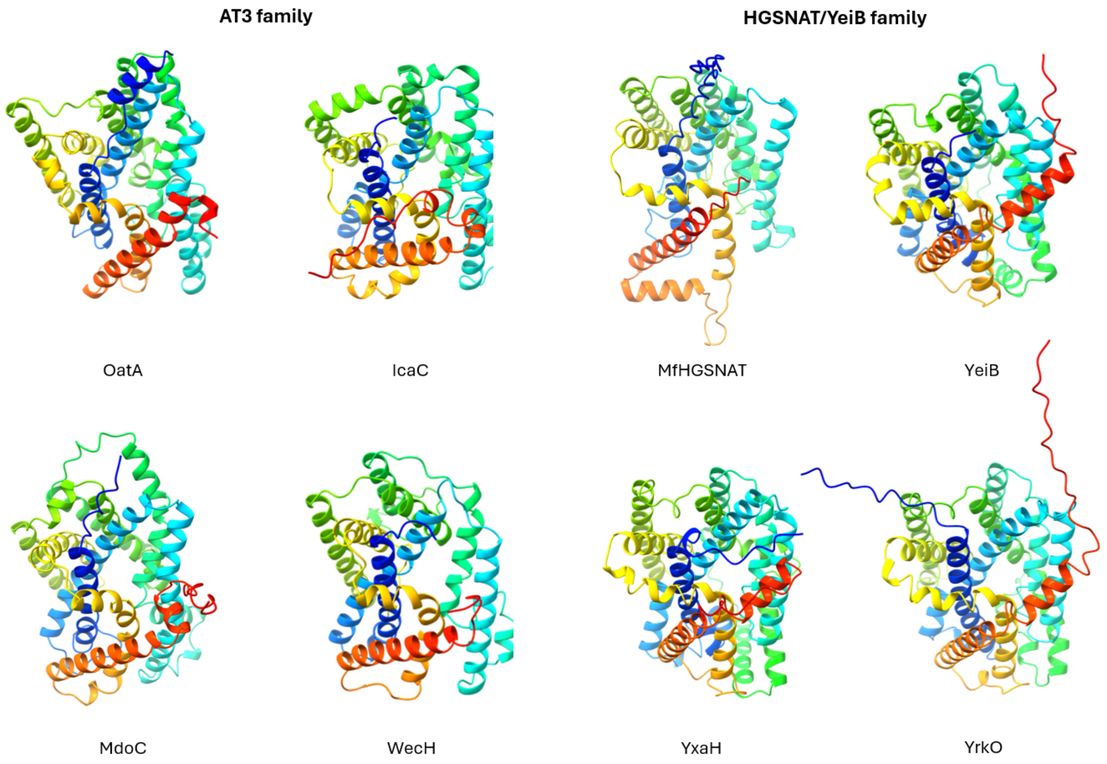


**Supplementary Fig.4. Conservation of a core 10-TMH TmAT protein fold within the two TDBC-defined families.** A cytosolic view of representative members of the AT3 and HGSNAT/YeiB families. Any additional domains and helices have been removed to illustrate the TmAT fold. Proteins are all coloured with rainbow colouring (N-terminus to C-terminus). All structures are Alphafold models. AT3 family representatives are OatA (Uniprot ID: Q2FV54), IcaC (Uniprot ID: Q5HKP7), MdoC (Uniprot ID: P75920) and WecH (Uniprot ID: P37669). HGSNAT/YeiB family representative members are MfHGSNAT (Uniprot ID: K2QAW2), YeiB (Uniprot ID: P254747), YxaH (Uniprot ID: P42107) and YrkO (Uniprot ID: P54442).

**Supplementary Table 1. The 32 proteins analysed within the TCDB-defined TmAT superfamily, along with the corresponding residues to which each structure was truncated.** These residues are inclusive. If a protein already contained 10TMH and was not truncated, it is marked as (uncut).

| Protein | Residues |
|---|---|
| AT3_Bac_S.paratyphi_OafB_A0A0H2WM30 | 1-343 |
| HGSNAT/YeiB_Arc_M. formicicum_K2QAW2 | 1-382 (uncut) |
| AT3_Bac_E.coli_WecH_P37669 | 1-331 (uncut) |
| HGSNAT/YeiB_Bac_E.coli_YeiB_P25747 | 1-385 (uncut) |
| HGSNAT/YeiB_Bac_B.subtilis_YrkO_P54442 | 1-405 (uncut) |
| AT3_Euk_D.melanogaster_Dropdead_Q8IR42 | 382-786 |
| AT3_Euk_M.musculus_Oacyl_Q8BML2 | 266-685 |
| AT3_Euk_P.amygdali_PaAT1_L0MXJ3 | 1-497 (uncut) |
| HGSNAT/YeiB_Bac_B.subtilis_YxaH_P42107 | 1-402 (uncut) |
| HGSNAT/YeiB_Euk_H.sapiens_HGSNAT_8jkv | 266-663 |
| AT3_Bac_S.epidermidis_IcaC_Q5HKP7 | 1-355 (uncut) |
| AT3_Bac_S.aureus_OatA_Q2FV54 | 1-368 (uncut) |
| AT3_Bac_E.coli_MdoC_P75920 | 1-385 (uncut) |
| HGSNAT/YeiB_Euk_T.sphaerospora_A0A4P9XPE3 | 68-472 |
| HGSNAT/YeiB_Bac_S.typhirium_E8XDM9 | 1-386 (uncut) |
| HGSNAT/YeiB_Euk_A.sinensis_A0A084WJN1 | 188-577 |
| HGSNAT/YeiB_ Euk_O.sativa_Q5WMX9 | 1-491 (uncut) |
| HGSNAT/YeiB_Euk_A.forsteri_A0A087QR62 | 159-551 |
| HGSNAT/YeiB_Euk_M.musculus_HGSNAT_Q3UDW8 | 257-656 |
| HGSNAT/YeiB_ Arc_H.larsenii_A0A1H7QXA1 | 1-364 (uncut) |
| HGSNAT/YeiB_ Arc_C.lokiarchaeum_A0A0F8XRW0 | 1-373 (uncut) |
| HGSNAT/YeiB_Arc_M.acetivorans_Q8TNU4 | 1-353 (uncut) |
| HGSNAT/YeiB_ Arc_H.lacisalsi_M0L9A7 | 1-432 (uncut) |
| HGSNAT/YeiB_Bac_B.acidopullulyticus_YAMY_P32819 | 1-346 (uncut) |
| AT3_ Arc_M.formicicum_A0A089ZV86 | 1-364 (uncut) |
| AT3_ Arc_M.nitroreducens_A0A062V6W2 | 1-379 (uncut) |

| | |
|---|---|
| AT3_ Arc_M.millerae_A0A0U2SIX7 | 1-349 (uncut) |
| AT3_ Arc_M.vacuolata_A0A0E3LGU3 | 1-245 (uncut) |
| AT3_ Arc_M.thermophila_A0A0E3H889 | 1-431 (uncut) |
| AT3_ Euk_C.lunatus_Clz18_A0A345BJP8 | 1-469 (uncut) |
| AT3_ Euk_C.elegans_NRF6_Q09225 | 372-822 |
| AT3_ Euk_C.elegans_RHY1_Q9XVW1 | 87-502 |

**Supplementary Table 2. DALI structural alignment statistics of AT3 and HGSNAT/YeiB family members when compared to HGSNAT (PDB ID: 8JKV).**

| Protein | Z score | RMSD | Aligned residues | Total no. of residues | Sequence ID % |
|---|---|---|---|---|---|
| HGSNAT/YeiB_Euk_M.musculus_HGSNAT_Q3UDW8 | 62.6 | 0.8 | 391 | 400 | 88 |
| HGSNAT/YeiB_Euk_A.forsteri_A0A087QR62 | 56.3 | 1.4 | 383 | 393 | 70 |
| HGSNAT/YeiB_Euk_A.sinensis_A0A084WJN1 | 55.9 | 1.3 | 384 | 390 | 45 |
| HGSNAT/YeiB_Euk_T.sphaerospora_A0A4P9XPE3 | 41.9 | 2.4 | 348 | 405 | 27 |
| HGSNAT/YeiB_Arc_M.formicicum_K2QAW2 | 40.7 | 2.1 | 335 | 382 | 29 |
| HGSNAT/YeiB_Euk_O.sativa_Q5WMX9 | 39.2 | 2.0 | 364 | 491 | 30 |
| HGSNAT/YeiB_Arc_C.lokiarchaeum_A0A0F8XRW0 | 26.2 | 3.1 | 301 | 373 | 16 |
| HGSNAT/YeiB_Arc_H.larsenii_A0A1H7QXA1 | 25.2 | 4.0 | 300 | 364 | 15 |
| HGSNAT/YeiB_Arc_M.acetivorans_Q8TNU4 | 25.1 | 3.4 | 298 | 353 | 19 |
| HGSNAT/YeiB_Bac_B.subtilis_YrkO_P54442 | 22.0 | 3.7 | 291 | 405 | 12 |
| HGSNAT/YeiB_Bac_B.subtilis_YxaH_P42107 | 20.9 | 3.9 | 282 | 402 | 15 |
| HGSNAT/YeiB_Bac_S.typhirium_E8XDM9 | 19.3 | 4.0 | 278 | 386 | 16 |
| HGSNAT/YeiB_Bac_E.coli_YeiB_P25747 | 19.3 | 3.9 | 280 | 385 | 17 |
| AT3_Arc_M.vacuolata_A0A0E3LGU3 | 18.9 | 4.1 | 280 | 345 | 14 |
| HGSNAT/YeiB_Arc_H.lacisalsi_M0L9A7 | 18.8 | 4.3 | 283 | 432 | 17 |
| HGSNAT/YeiB_Bac_B.acidopullulyticus_YAMY_P32819 | 18.3 | 3.8 | 263 | 346 | 16 |
| AT3_Bac_S.epidermidis_IcaC_Q5HKP7 | 18.2 | 4.7 | 286 | 355 | 10 |
| AT3_Euk_M.musculus_Oacyl_Q8BML2 | 16.3 | 4.7 | 291 | 420 | 10 |
| AT3_Euk_C.elegans_RHY1_Q9XVW1 | 15.7 | 5.0 | 285 | 416 | 13 |
| AT3_Arc_M.thermophila_A0A0E3H889 | 15.6 | 4.4 | 272 | 431 | 14 |
| AT3_Euk_D.melanogaster_Dropdead_Q8IR42 | 15.1 | 4.8 | 276 | 405 | 13 |
| AT3_Bac_E.coli_MdoC_P75920 | 14.9 | 4.9 | 289 | 385 | 11 |
| AT3_Euk_C.elegans_NRF6_Q09225 | 14.9 | 4.7 | 293 | 451 | 13 |
| AT3_Arc_M.millerae_A0A0U2SIX7 | 14.5 | 4.7 | 261 | 349 | 11 |
| AT3_Bac_S.aureus_OatA_Q2FV54 | 14.0 | 4.7 | 250 | 368 | 16 |
| AT3_Arc_M.nitroreducens_A0A062V6W2 | 13.8 | 4.5 | 271 | 379 | 13 |
| AT3_Bac_E.coli_WecH_P37669 | 13.5 | 5.7 | 250 | 331 | 13 |
| AT3_Arc_M.formicicum_A0A089ZV86 | 13.4 | 4.4 | 248 | 364 | 10 |
| AT3_Euk_P.amygdali_PaAT1_L0MXJ3 | 13.3 | 4.6 | 266 | 497 | 15 |
| AT3_Bac_S.paratyphi_OafB_A0A0H2WM30 | 13.2 | 5.0 | 250 | 343 | 12 |
| AT3_Euk_C.lunatus_Clz18_A0A345BJP8 | 13.2 | 4.5 | 258 | 469 | 12 |

**Supplementary Table 3. The 9 Pfam families that constitute the Pfam Acyl_transf_3 Clan (CL0316).** Pfam families where members are included in TCDB are mentioned with their TC code.

| Pfam family | Name | TCDB | TmAT | Example |
|---|---|---|---|---|
| PF01757 | Acyl_transf_3 (AT3) | 9.B.97 | Yes | *S.* Typhimurium OafB |
| PF04235 | DUF418 | 9.B.169 | Yes | *E. coli* YeiB |
| PF07768 | HGSNAT-cat | 9.B.169 | Yes | Human HGSNAT structures |
| PF05857 | TraX[1] | N/A | Yes* | *E. coli* TraX (plasmid bourne) |
| PF07779 | Cas1_AcylT | N/A | Yes | *Cryptococcus neoformans* Cas1p |
| PF10129 | OpgC-C | N/A | Yes | *Ricinus communis* OpgC |
| PF11318 | DUF3120[2] | N/A | No | *Synechocystis* sp. Slr1990 |
| PF12291 | DUF3623[3] | N/A | No | *R. capsulatus* RCAP_rcc00656 |
| PF16401 | DUF5009[4] | 9.B.169 | Yes | *B. thetaiotaomicron* BT_0446 |

[1] TraX has 9 TMH in the fold, missing the usual TM10. Also, TM9 is not broken. Some of the TraX proteins included in this analysis do have 10TM helices (Uniprot ID: A0A653CKF7 and P22709) but the extra helix comes before the TmAT-defined TM1. Additionally, another couple of TraX proteins have a short TM1 (Uniprot ID: A0A497P6U4 and A0A077ZID6). There is a conserved $Kx_9DH$ motif in what would be TM1 and other helices provide conserved R residues on the potential acyl-CoA binding face, so there are certainly features that are conserved. Given its known biological function in acetylation, we support the hypothesis that this is a divergent member of the superfamily. Its divergence is perhaps related to its acceptor substrate which is thought to be an Alanine residue in the F-pilin protein and not a sugar[8,9].

[2]DUF3120 are shorter proteins than typical TmAT proteins, at around 200 amino acids and contain 6 TMH. Foldseek does not find significant matches to any known TmAT proteins, nor in fact, any confident matches to any other known protein fold.

[3] DUF3623 are around 280 amino acids, so shorter than regular TmAT proteins and are predicted to have 7 TMH. As for DUF3120, Foldseek does not find significant matches to any known TmAT proteins, nor in fact any confident matches to any other known fold.

[4]DUF5009 is similar to HGSNAT, although it has two sets of two additional TM helices on either side of the protein. It is suggested in TCDB to be part of the HGSNAT/YeiB family (9.B.169).

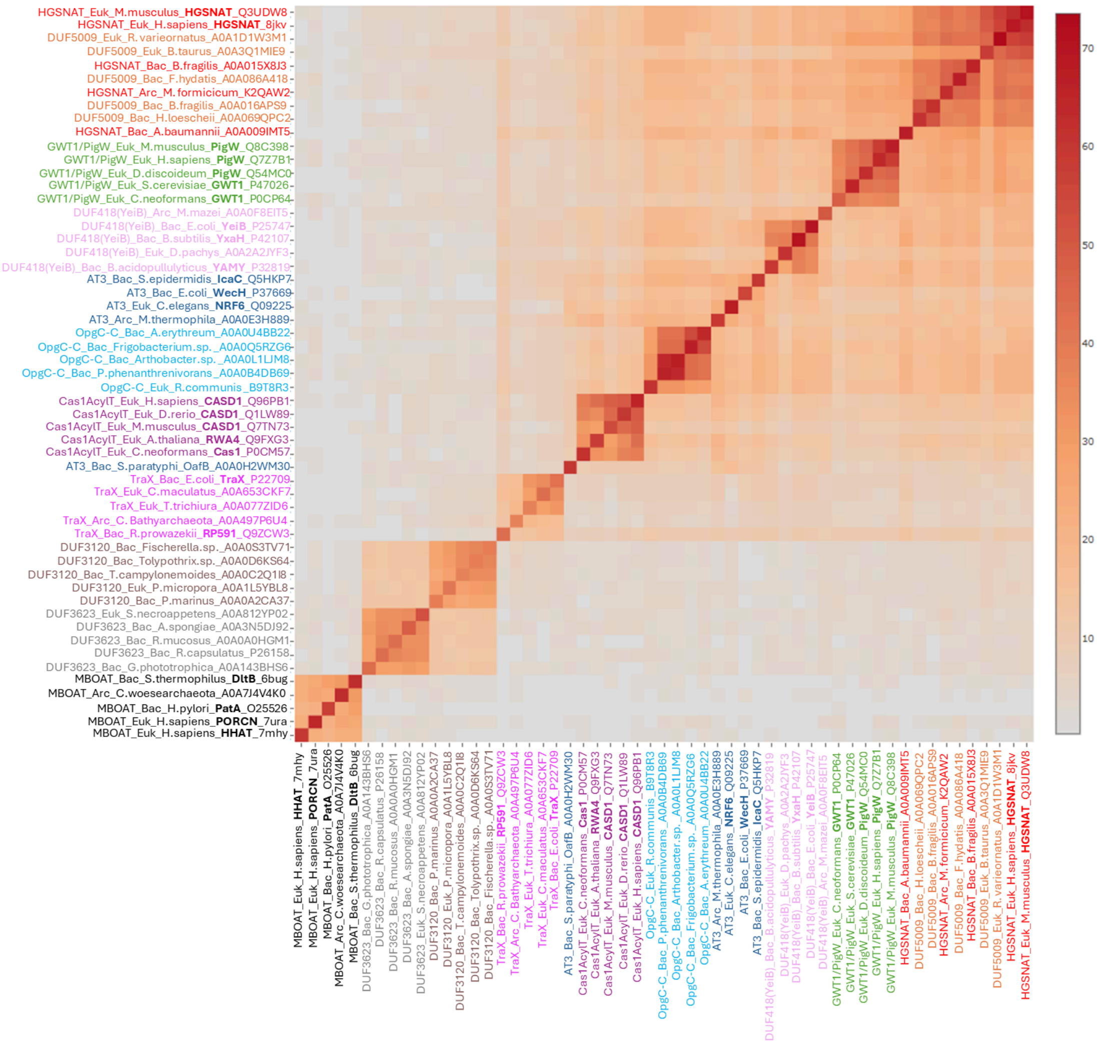


**Supplementary Fig.5. Heat map of Z-scores of a DALI all against all alignment of select members of the Pfam Acyl_transf_3 Clan (CL0316), GWT1/PIG-W family and MBOAT superfamily.** Z scores of the 10 families and 1 superfamily investigated with 5 representative members in each including members from archaea (Arc), bacteria (Bac) and eukaryota (Euk) are displayed. Members are coloured according to family - AT3 in dark blue, HGSNAT/YeiB in red, DUF5009 in orange, GWT1/PIG-W in green, DUF418 in light pink, OpgC-C in light blue, Cas1_AcylT in maroon, TraX in purple, DUF3120 in brown, DUF3623 in grey and MBOAT in black. Note that three proteins that were previously annotated as DUF5009 have now been annotated as HGSNAT or are unlabelled (see Methods). The average Z score between MBOAT and every protein included in this alignment serves as a negative control for structural similarity (the average Z score is 1.2). The average Z-score between all families (excluding those within the families) and excluding DUF3120, DUF3623, and MBOAT is 14.1, confirming significant structural similarity between AT3, HGSNAT/YeiB, DUF5009, GWT1/PIG-W, DUF418, OpgC-C, Cas1_AcylT and TraX families.

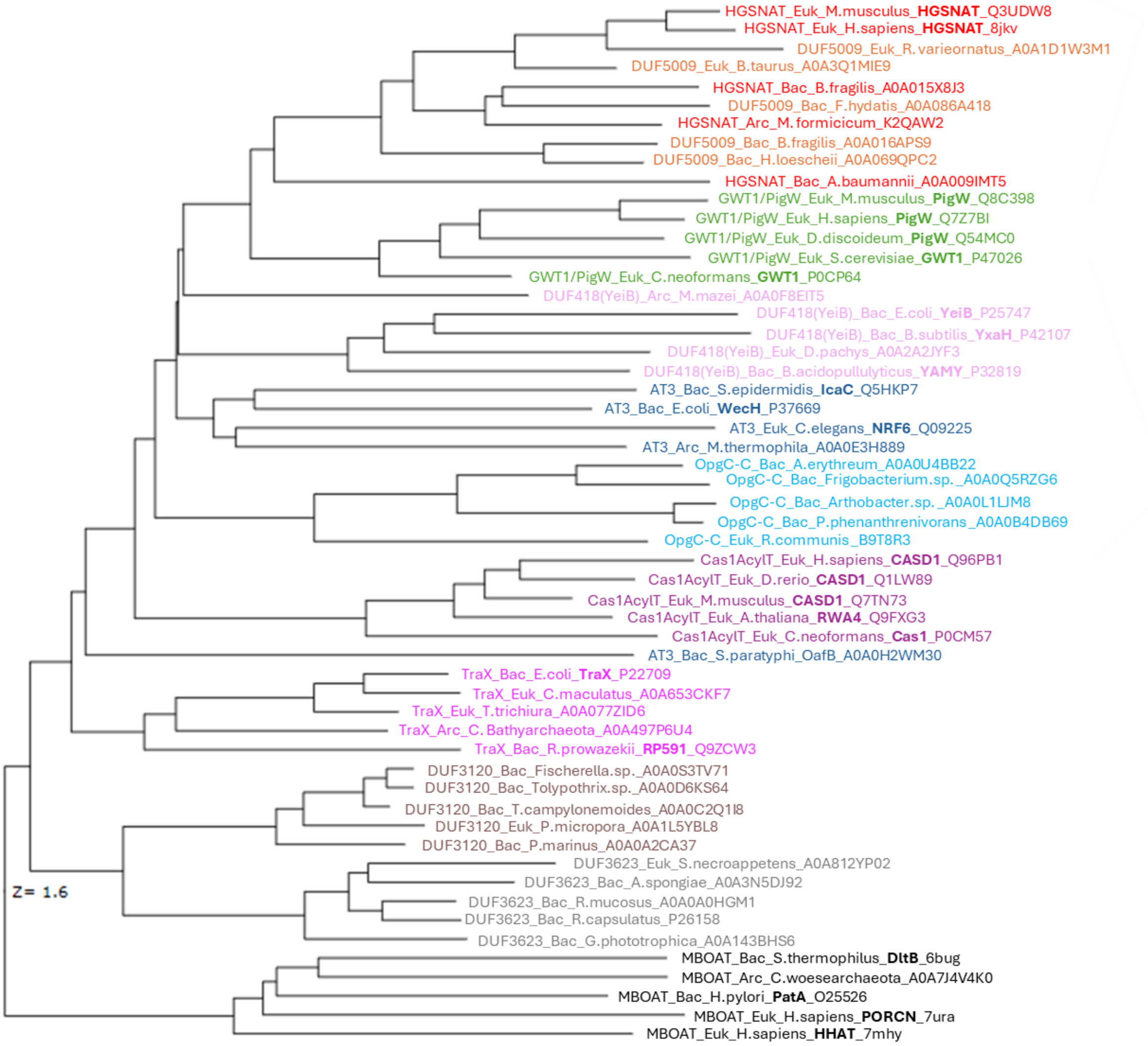


**Supplementary Fig.6. Structural dendrogram of select members of the Pfam Acyl_transf_3 Clan (CL0316), GWT1/PIG-W family and MBOAT superfamily.** Each group contains 5 representative proteins including members from archaea (Arc), bacteria (Bac) and eukaryota (Euk) where possible. The structural dendrogram shows that AT3, HGSNAT/YeiB, DUF5009, GWT1/PIG-W, DUF418, OpgC-C, Cas1_AcylT and TraX families are structurally related. It also highlights that the MBOAT superfamily is not structurally related to the other investigated families (Z score of 1.4). Members are coloured according to family - AT3 in dark blue, HGSNAT/YeiB in red, DUF5009 in orange, GWT1/PIG-W in green, DUF418 in light pink, OpgC-C in light blue, Cas1_AcylT in maroon, TraX in purple, DUF3120 in brown, DUF3623 in grey and MBOAT in black. Note that three proteins that were previously annotated as DUF5009 have now been annotated as HGSNAT or are unlabelled (see Methods).

**Supplementary Table 4. DALI structural alignment statistics of the Pfam Acyl_transf_3 Clan (CL0316), GWT1/PIG-W family and MBOAT superfamily members when compared to HGSNAT (PDB ID: 8JKV).** Proteins with Z scores below 2 are not shown. Note that three proteins that were previously annotated as DUF5009 have now been annotated as HGSNAT or are unlabelled (see Methods).

| Protein | Z score | RMSD | Aligned residues | Total no. of residues | Sequence ID % |
|---|---|---|---|---|---|
| HGSNAT_Euk_H.sapiens_HGSNAT_8jkv | 69.0 | 0.0 | 527 | 527 | 100 |
| HGSNAT_Euk_M.musculus_HGSNAT_Q3UDW8 | 62.6 | 0.9 | 527 | 656 | 84 |
| DUF5009_Euk_R. varieornatus_A0A1D1W3M1 | 56.0 | 1.4 | 508 | 1195 | 44 |
| DUF5009_Euk_B.taurus_A0A3Q1MIE9 | 49.0 | 1.4 | 484 | 690 | 81 |
| HGSNAT_Bac_B.fragilis_A0A015X8J3 | 41.0 | 2.1 | 335 | 387 | 30 |
| HGSNAT_Arc_M. formicicum_K2QAW2 | 40.6 | 2.1 | 335 | 382 | 29 |
| DUF5009_Bac_F.hydatis_A0A086A418 | 39.1 | 2.5 | 340 | 423 | 31 |
| DUF5009_Bac_H.loescheii_A0A069QPC2 | 36.6 | 2.6 | 332 | 409 | 23 |
| DUF5009_Bac_B.fragilis_A0A016APS9 | 35.8 | 2.7 | 328 | 375 | 28 |
| HGSNAT_Bac_A.baumannii_A0A009IMT5 | 26.8 | 3.0 | 298 | 381 | 16 |
| GWT1/PigW_Euk_M.musculus_PigW_Q8C398 | 24.2 | 3.3 | 326 | 503 | 13 |
| GWT1/PigW_Euk_S.cerevisiae_GWT1_P47026 | 23.8 | 3.5 | 328 | 490 | 15 |
| GWT1/PigW_Euk_D.discoideum_PigW_Q54MC0 | 23.4 | 3.5 | 330 | 492 | 11 |
| GWT1/PigW_Euk_H.sapiens_PigW_Q7Z7B1 | 23.2 | 3.3 | 325 | 504 | 14 |
| DUF418(YeiB)_Bac_B.subtilis_YxaH_P42107 | 21.0 | 3.9 | 283 | 402 | 15 |
| OpgC-C_Euk_R.communis _B9T8R3 | 20.6 | 4.5 | 293 | 369 | 14 |
| OpgC-C_Bac_Arthobacter.sp. _A0A0L1LJM8 | 20.4 | 4.0 | 313 | 795 | 12 |
| OpgC-C_Bac_P.phenanthrenivorans _ A0A0B4DB69 | 20.2 | 4.0 | 327 | 831 | 12 |
| DUF418(YeiB)_Arc_M.mazei_A0A0F8EIT5 | 19.7 | 3.9 | 286 | 340 | 16 |
| OpgC-C_Bac_Frigobacterium.sp. _A0A0Q5RZG6 | 19.6 | 4.1 | 293 | 824 | 11 |
| GWT1/PigW_Euk_C.neoformans_GWT1_P0CP64 | 19.2 | 3.9 | 321 | 598 | 12 |
| DUF418(YeiB)_Bac_E.coli_YeiB_P25747 | 18.9 | 4.1 | 283 | 385 | 14 |
| OpgC-C_Bac_A.erythreum_A0A0U4BB22 | 18.7 | 5.3 | 291 | 811 | 12 |
| DUF418(YeiB)_Bac_B.acidopullulyticus_YAMY_ P32819 | 18.3 | 3.8 | 264 | 346 | 16 |
| AT3_Bac_S.epidermidis_IcaC_Q5HKP7 | 17.6 | 4.6 | 290 | 355 | 10 |
| AT3_Arc_M.thermophila_A0A0E3H889 | 15.7 | 4.4 | 274 | 431 | 14 |
| DUF418(YeiB)_Euk_D.pachys_A0A2A2JYF3 | 15.4 | 4.7 | 269 | 375 | 12 |
| Cas1AcylT_Euk_H.sapiens_CASD1_Q96PB1 | 15.2 | 6.2 | 300 | 797 | 11 |

| | | | | | |
|---|---|---|---|---|---|
| AT3_Euk_C.elegans_NRF6_Q09225 | 15.0 | 6.0 | 308 | 822 | 11 |
| Cas1AcylT_Euk_D.rerio_CASD1_Q1LW89 | 14.5 | 6.0 | 284 | 781 | 10 |
| Cas1AcylT_Euk_A.thaliana_RWA4_Q9FXG3 | 13.6 | 4.6 | 284 | 540 | 10 |
| AT3_Bac_E.coli_WecH_P37669 | 13.3 | 4.9 | 249 | 331 | 14 |
| Cas1AcylT_Euk_C.neoformans_Cas1_P0CM57 | 12.3 | 5.2 | 299 | 960 | 8 |
| Cas1AcylT_Euk_M.musculus_CASD1_Q7TN73 | 11.8 | 6.2 | 298 | 797 | 12 |
| TraX_Bac_R.prowazekii_RP591_Q9ZCW3 | 9.7 | 5.8 | 189 | 248 | 13 |
| AT3_Bac_S.paratyphi_OafB_A0A0H2WM30 | 7.8 | 4.7 | 243 | 640 | 12 |
| TraX_Bac_E.coli_TraX_P22709 | 7.6 | 4.0 | 168 | 248 | 13 |
| DUF3120_Euk_P.micropora_A0A1L5YBL8 | 6.5 | 6.4 | 156 | 244 | 6 |
| TraX_Euk_C.maculatus_A0A653CKF7 | 6.1 | 4.6 | 165 | 245 | 10 |
| TraX_Arc_C. Bathyarchaeota_A0A497P6U4 | 5.8 | 4.6 | 165 | 209 | 12 |
| DUF3120_Bac_Fischerella.sp._A0A0S3TV71 | 5.7 | 4.8 | 158 | 215 | 7 |
| DUF3120_Bac_T.campylonemoides_A0A0C2Q1I8 | 5.6 | 9.6 | 184 | 246 | 6 |
| DUF3120_Bac_Tolypothrix.sp._A0A0D6KS64 | 5.5 | 5.1 | 145 | 259 | 8 |
| DUF3120_Bac_P.marinus_A0A0A2CA37 | 5.3 | 5.6 | 152 | 212 | 9 |
| TraX_Euk_T.trichiura_A0A077ZID6 | 5.1 | 4.7 | 151 | 205 | 11 |
| DUF3623_Bac_A.spongiae_A0A3N5DJ92 | 5.0 | 5.9 | 131 | 579 | 6 |
| DUF3623_Euk_S.necroappetens_A0A812YP02 | 4.7 | 7.0 | 175 | 824 | 9 |
| DUF3623_Bac_R.capsulatus_P26158 | 4.5 | 4.6 | 151 | 274 | 9 |
| DUF3623_Bac_G.phototrophica_A0A143BHS6 | 4.4 | 5.5 | 163 | 301 | 7 |
| DUF3623_Bac_R.mucosus_A0A0A0HGM1 | 3.7 | 5.5 | 144 | 270 | 6 |
| MBOAT_Euk_H.sapiens_PORCN_7ura | 3.4 | 3.6 | 58 | 432 | 5 |
| MBOAT_Bac_S.thermophilus_DltB_6bug | 3.2 | 6.0 | 122 | 414 | 8 |

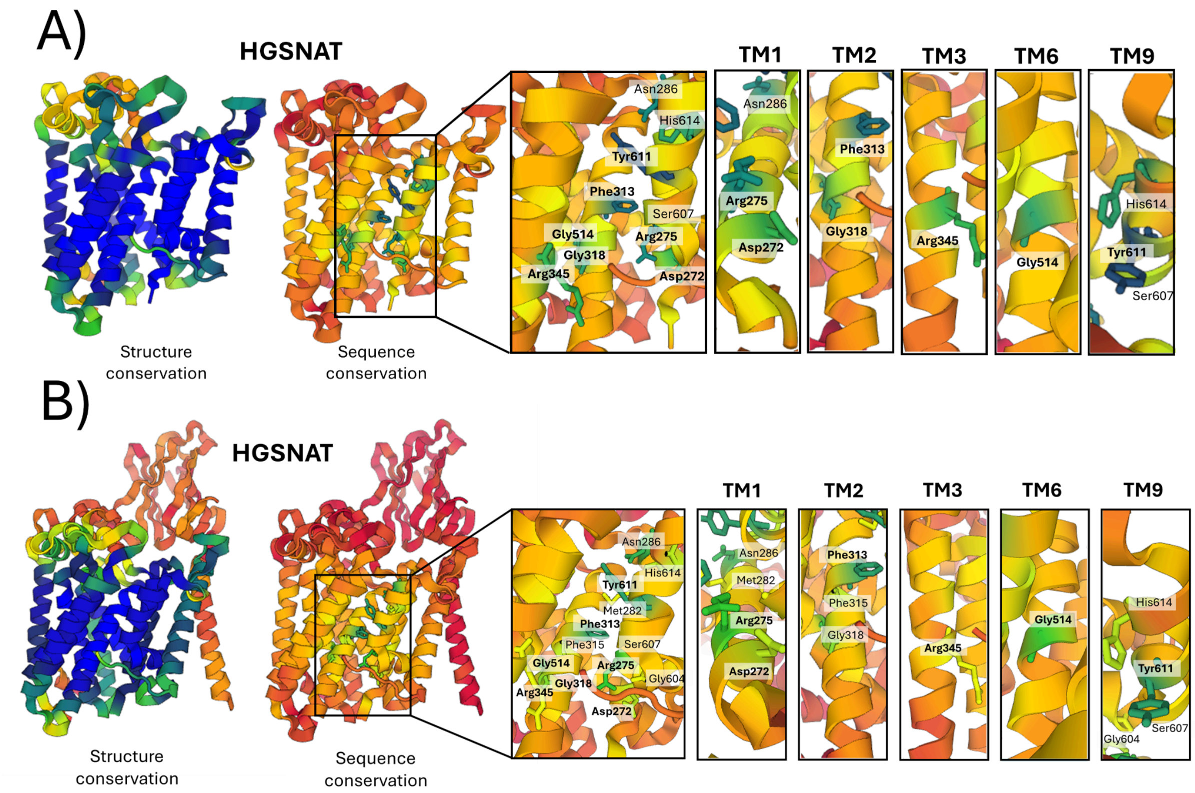


**Supplementary Fig.7. Protein structure and amino acid position conservation between select members containing a TmAT domain.** A) Conserved amino acid positions across the TCDB-defined TmAT superfamily (HGSNAT/YeiB and AT3 family members that are truncated to only include the core 10 TM TmAT domain) overlaid on the TmAT domain of human HGSNAT (PDB: 8JKV) structure generated using DALI. B)  Conserved amino acid positions across the TmAT superfamily (HGSNAT, DUF418/YeiB, OpgC-C, AT3, Cas1_AcylT, PigW/GWT1, and TraX) overlaid on the human HGSNAT (PDB: 8JKV) structure generated using DALI. The left models show structural conservation while the right shows sequence conservation. A colour gradient from blue to red is used to indicate conservation with blue indicating the most conserved features and red indicating the least conserved features. Panels on the right-hand side highlight the conservation of residues in TMH 1-3, 6 and 9 (using TmAT helical numbering). Highly conserved residues are highlighted in bold.

**Supplementary Table 5.** The 7 conserved residues of the TmAT superfamily (along with the partially conserved TMH9-Ser) are listed with their corresponding residue numbers and respective helices in both HGSNAT and OafB.

| **TMH-residue** | **HGSNAT** | **OafB** |
|---|---|---|
| TMH1-Asp | 272 | 11 |
| TMH1-Arg | 275 | 14 |
| TMH2-Phe | 313 | 41 |
| TMH2-Gly | 318 | 46 |
| TMH3-Arg | 345 | 71 |
| TMH6-Gly | 514 | 207 |
| TMH9-Tyr | 611 | 292 |
| TMH9-Ser | 607 | 288 |

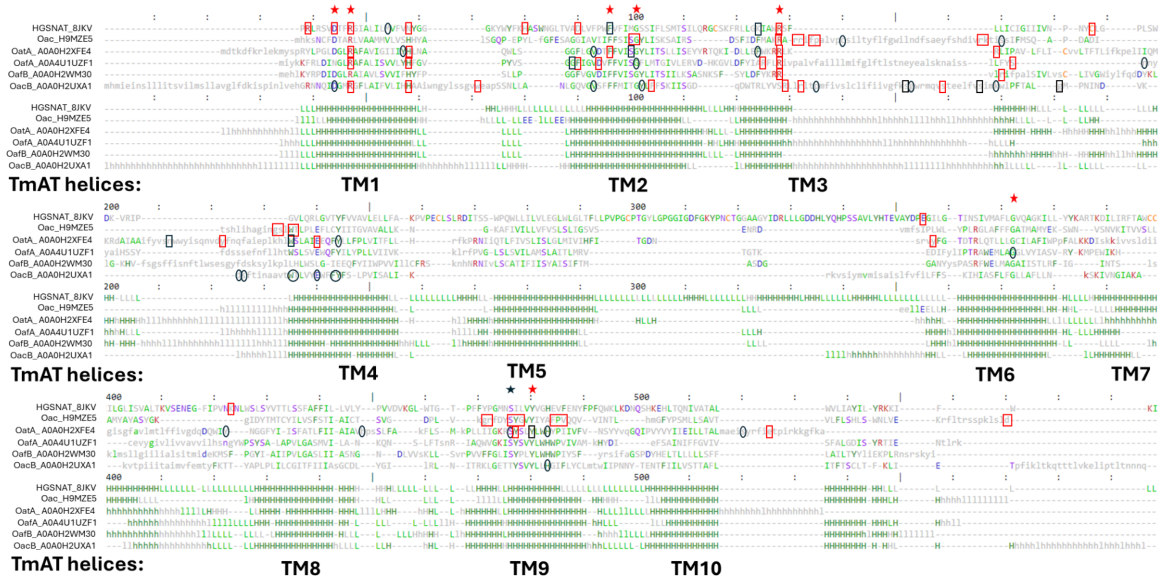


**Supplementary Fig.8. Structure-based sequence alignment highlighting amino acid residues in HGSNAT and select AT3 members that have been functionally analysed by mutagenesis.** HGSNAT (PDB ID: 8JKV), Oac (Uniprot ID: H9MZE5), OatA (Uniprot ID: A0A0H2XFE4), OafA (Uniprot ID: A0A4U1UZF1), OafB (Uniprot ID: A0A0H2WM30), and OacB (Uniprot ID: A0A0H2UXA1) were truncated to include only their 10TM TmAT domains. Mutated residues, highlighted by black boxes, have been identified as affecting protein function. A red box marks residues deemed critical to protein activity, either based on supporting literature[10–13] or when mutations result in at least a 50% reduction in activity. Residues that are circled in black were also mutated but were found to be non-critical. In cases where multiple residues are boxed together, it indicates that they were mutated simultaneously. The 7 residues we identify as being conserved across the TmAT superfamily are marked with a red star above. The additional partially conserved residue, $Ser607_{HGSNAT}$, that is discussed in the text is marked with a black star. Protein secondary structure is annotated as H - Helix, L - Coil, and E -Sheet.

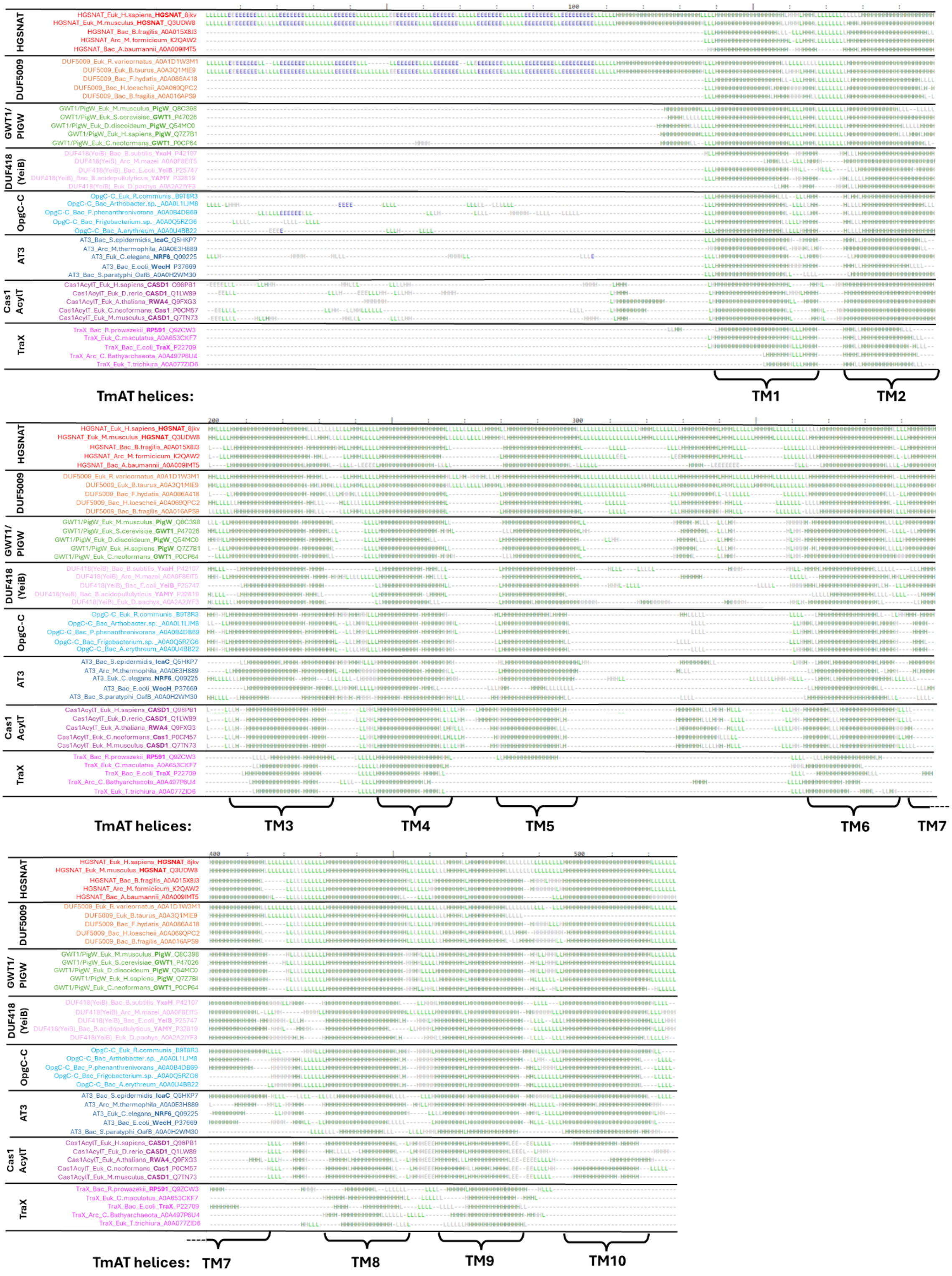


**Supplementary Fig.9. Secondary structure alignment of the TmAT superfamily members.** Secondary structure alignment (corresponding to **Fig.3**) compared to HGSNAT 8JKV) produced in DALI showing diverse representatives of Pfam Acyl_transf_3 Clan (CL0316) members containing the TmAT domain and the additional PigW/GWT1 family found by a DALI search. Regions with no structural/sequence homology are not expanded, so only structurally equivalent positions to HGSNAT (8JKV) are shown. It is important to note that some of the proteins appear to not have a TM5 or TM7, but this is due to poor structural alignment with other TmAT members and not because they lack a TM5 or 7. Protein secondary structure is annotated as H - Helix, L - Coil, and E -Sheet.

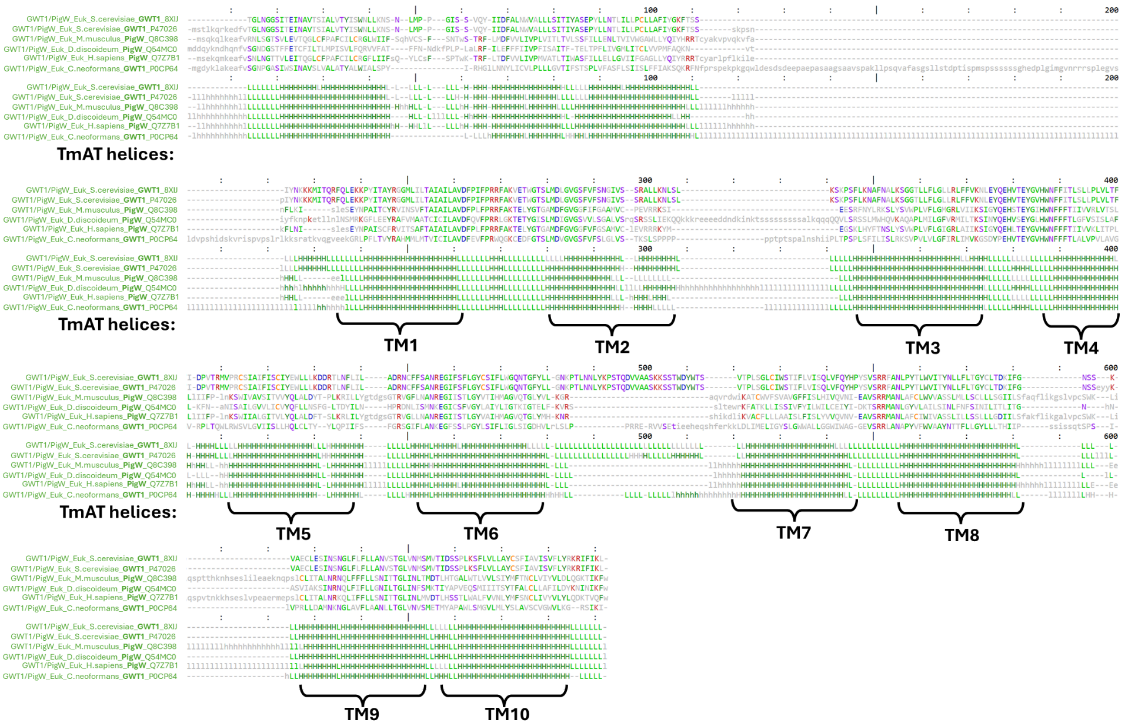


**Supplementary Fig.10. Structure-based multiple sequence alignment of GWT1/PigW proteins used in the analysis compared to the recently solved structure of GWT1 (PDB ID: 8XIJ).** The alignments were produced using DALI and GWT1 (PDB: 8XIJ) as the reference structure. This alignment demonstrates that the sequence and structure of the resolved GWT1 closely resemble the AlphaFold GWT1/PigW structures, particularly for the GWT1 protein from *Saccharomyces cerevisiae* (Uniprot ID: P47026). Regions lacking any sequence/structural similarity are not expanded, so only structurally equivalent positions to GWT1 (PDB: 8XIJ) are shown. The most common amino acid at any given position is coloured according to aromaticity (dark green)(Phe, His, Trp, Tyr), positive charge (red)(Arg, Lys), negative charge (blue)(Glu, Asp), polar uncharged side chains (purple) (Ser, Thr, Gln, Asn), hydrophobic side chains that are not aromatic (Gly, Leu, Ile, Val, Pro, Met, Ala), and sulfur-containing amino acid (orange) (Cys). Protein secondary structure is annotated as H - Helix, L - Coil, and E -Sheet.

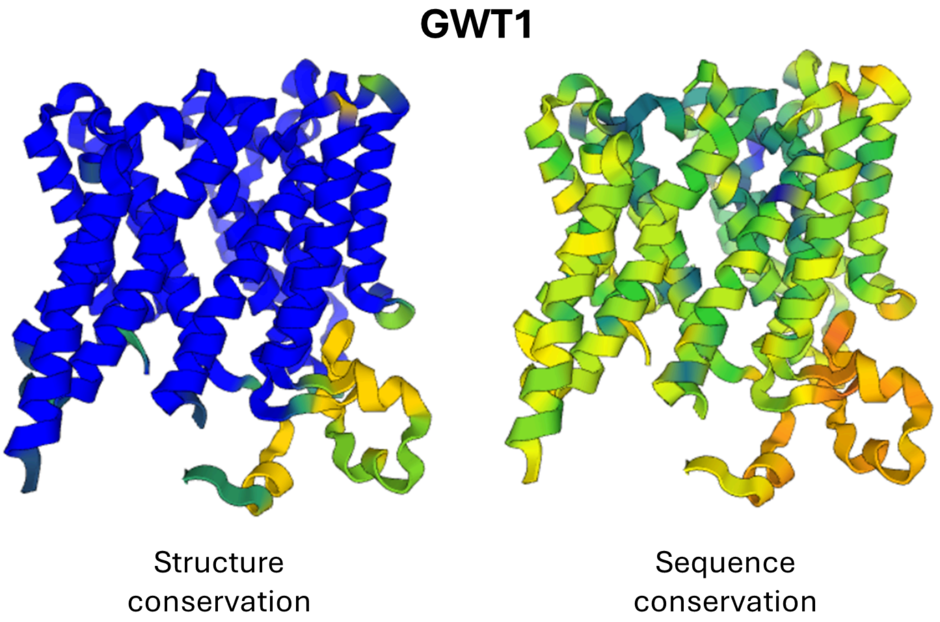


**Supplementary Fig.11. Protein structure and amino acid position conservation between GWT1/PigW proteins used in the analysis and the recently solved structure of GWT1 (PDB ID: 8XIJ)** Conserved amino acid positions across the GWT1 superfamily (Uniprot IDs: P47026, Q8C398, Q54MC0, Q7Z7B1, P0CP64) overlaid on the GWT1 (PDB: 8XIJ) structure generated using DALI. The left models show structural conservation while the right shows sequence conservation. A colour gradient from blue to red is used to indicate conservation with blue indicating the most conserved features and red indicating the least conserved features. The structural and sequence conservation demonstrate that the solved structure of GWT1 closely resembles the AlphaFold GWT1/PigW structures used in this analysis.

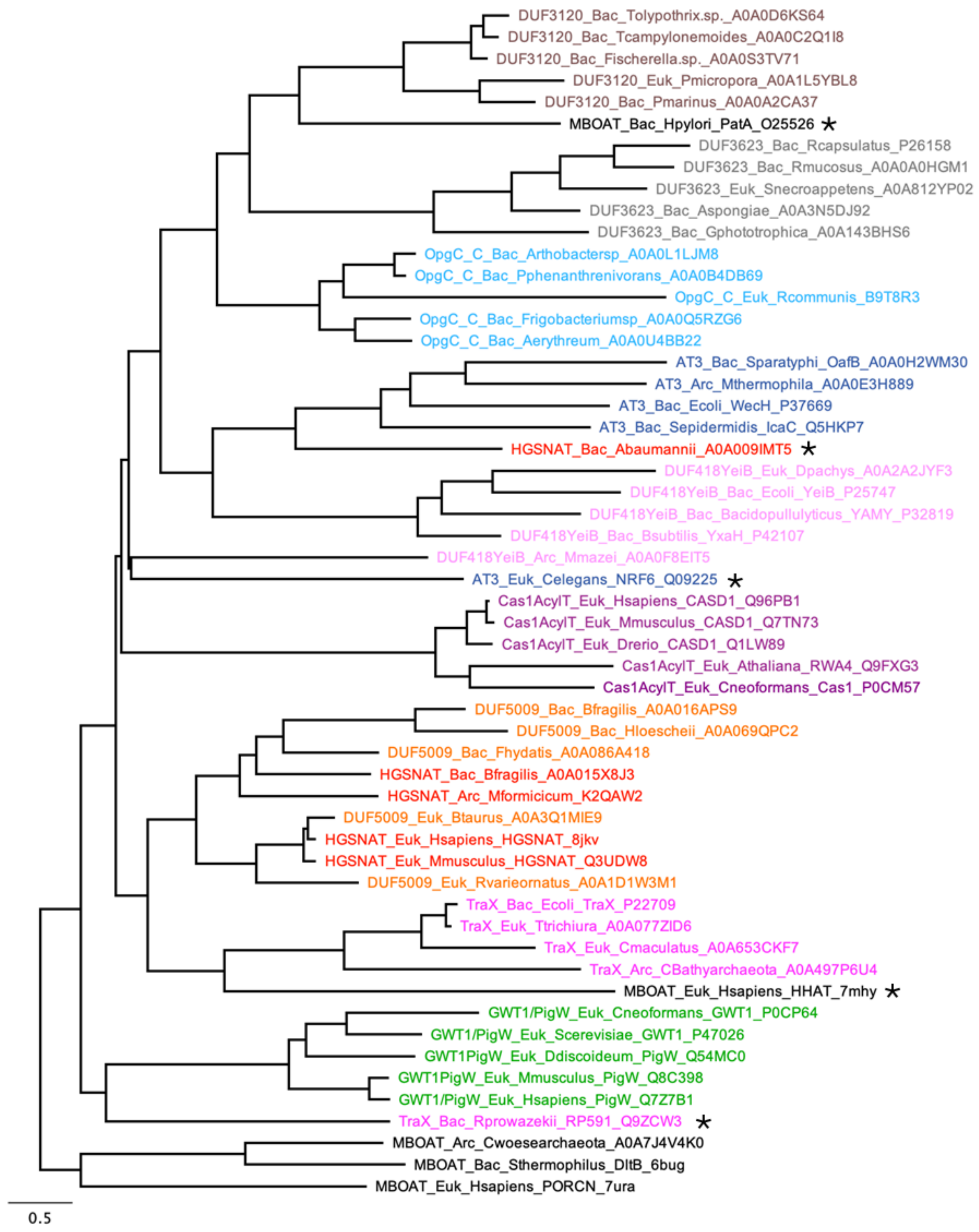


**Supplementary Figure 12. Maximum likelihood phylogenetic tree of select members of the Pfam Acyl_transf_3 Clan (CL0316), GWT1/PIG-W family and MBOAT superfamily.** Refining the TmAT superfamily remains challenging when solely relying on sequence alignments, given the low sequence similarity between these proteins. This phylogenetic tree positions the DUF3120, DUF3623, and MBOAT proteins as more closely related to the other protein families, in contrast to the structural dendrogram (***Supplementary Fig. 6***), which places them as the least related to the other proteins. Asterisks highlight proteins that are inaccurately distanced from their other family members. This demonstrates the limitations of relying solely on sequence alignments, as opposed to structural analysis, for protein classification.
